# What doesn’t kill you makes you stronger? Effects of paternal age at conception on fathers and sons

**DOI:** 10.1101/2023.12.20.572572

**Authors:** Krish Sanghvi, Tommaso Pizzari, Irem Sepil

## Abstract

Advancing male age is often hypothesised to reduce both male fertility and offspring quality through reproductive senescence. However, the effects of advancing male age on reproductive output and offspring quality need not always be deleterious. For example, older fathers might compensate for reproductive senescence by terminally investing in reproduction. Similarly, males that survive to reproduce at an old age, might carry alleles that confer high viability (viability selection) which are then inherited by offspring, or might have high reproductive potential (selective disappearance). Differentiating these mechanisms requires an integrated experimental study of paternal survival and reproductive performance, as well as offspring quality, which is currently lacking. Using *Drosophila melanogaster*, we test the effects of paternal age at conception (PAC) on paternal survival and reproductive success, and the lifespans of sons. We discover that mating at an old age is temporarily linked with decreased future male survival, suggesting that mating-induced mortality is possibly due to old fathers being frail. We also find a quadratic reproductive ageing pattern, with an onset of senescence in late-life. We discover no evidence for terminal investment, and instead discover positive covariances between a father’s lifespan and his probability of siring offspring, for older PAC groups. Lastly, we show that sons born to older fathers live longer than those born to younger fathers, due to viability selection. Collectively, our results suggest that effects of advancing PAC need not be deleterious for fathers or offspring, and can increase fitness if older fathers produce more viable offspring.

**Lay summary:** It’s often assumed that old fathers have fewer or lower-quality offspring than young fathers. However, old fathers have on average, usually survived to and reproduced at an older age than young fathers, which might be signals of high quality of old fathers, and benefit offspring. These opposing predictions have rarely been tested, and it’s unclear what their combined influence is, in determining the father’s and offspring’s fitness. Using fruit flies, we explored how old paternal age affects a father’s reproduction and his sons’ lifespans. We find evidence for age-related reproductive decline in fathers. However, we also discover that fathers who mate at an older age live longer than fathers who mate at a younger age, and subsequently, the sons of older fathers live longer than sons of younger fathers. We suggest that the relationship between paternal age, paternal reproduction and lifespan, and offspring quality, is more complex than previously assumed, and that old fathers might provide benefits to their offspring that future studies need to consider.

**Teaser text:** Is an old dad really that bad? While old fathers might in some cases produce fewer offspring or offspring of worse quality than young fathers, they have also survived to an older age, thus might be of higher quality. We test how reproducing at an old age affects a father and his son’s fitness. We show that old fathers show evidence of reproductive senescence, but, surprisingly, produce sons that are longer lived than the sons of young fathers. This effect is driven largely by old fathers themselves living longer than young fathers, thus possibly passing alleles conferring higher viability to their sons. Our research is important because it reveals specific mechanisms that drive multifaceted effects of age on fitness.

## Introduction

Reproductive senescence is the age-dependent decline in the reproductive output of organisms (Monaghan et al, 2008; Rose and Charlesworth, 1980). This leads old individuals to have lower gamete quality and quantity (Dean et al, 2010; Gasparini et al, 2010), thus lower fecundity and fertility (Naciri et al, 2022; Sepil et al, 2020), and produce fewer offspring, than young individuals. For instance, old males across different species often have smaller ejaculate sizes (Cornwallis et al, 2014; Sanghvi et al, 2023), poorer quality sperm (Gasparini et al, 2019; Johnson et al, 2015), and lower abundance of seminal fluid proteins (Fricke et al, 2023), than young males. A growing body of literature however, challenges these patterns by documenting that advancing male age does not necessarily result in male reproductive senescence (Aich et al, 2022; Baudisch and Stott, 2019; Brooks and Kemp, 2001; Cooper et al, 2020, 2021; Finch, 2009; Forslund and Part, 1995; Heinze et al, 2018; Johnson and Gemmell, 2012; Jones et al, 2014; Jones and Vaupel, 2017; Lee and Chu, 2023; Moullec et al, 2023; Sandfoss et al, 2023; Segami et al, 2021; Vega-Trejo et al, 2019). In some cases, advancing age may even be associated with increased reproductive output (e.g. Avent et al, 2008; Girndt et al, 2019; Lifjeld et al, 2022; Prathibha et al, 2011; Sanghvi et al, 2023; Santhosh and Krishna, 2013; Verspoor et al, 2015).

Several mechanisms explain why old males might have a reproductive output similar to-, or higher than, young males. Individuals of low reproductive output may be more likely to die (i.e. selective disappearance) with increasing age (Bouwhuis et al, 2009; Hamalainen et al, 2014; Sanghvi et al, 2022; Sultanova et al, 2023), thus masking male reproductive senescence. Here, males who survive to an old age represent a non-random sample of high-quality individuals. Alternatively, according to the terminal investment hypothesis, old males might allocate proportionally more resources (e.g. ejaculate expenditure) to a single current reproductive event than young males (Farchmin et al, 2020; Moullec et al, 2023). This is because as future survival prospects of organisms decline, individuals might invest more in current than future reproductive opportunities (Creighton et al, 2009; Duffield et al, 2018; Froy et al, 2013; Part et al, 1992; Velando et al, 2006).

In addition to impacting a male’s own reproductive output, reproductive senescence in males can also influence the phenotypes of the offspring via paternal age effects (Priest et al, 2007; Schroeder et al, 2015). For instance, offspring sired by old fathers are reported to have poorer development (Janecka et al, 2017; Preston et al, 2015), early-life performance (Fay et al, 2016), and reproductive output (Arslan, 2017; Nystrand and Dowling, 2014; Vuarin et al, 2021), than offspring of young fathers. Notably, offspring born to old fathers often have shorter lifespans than those born to young fathers (Crow, 2003; Monaghan et al, 2020; Sharma et al, 2015), a phenomenon known as the ‘Lansing effect’ (Ivimey-Cook et al, 2023; Lansing, 1947). While these paternal age effects might be caused by age-dependent changes in paternal care (Benowitz et al, 2013; Cope et al, 2022), in species without care, these effects likely occur via age-dependent deterioration in ejaculates (Monaghan and Metcalfe, 2019).

In contrast to negative effects of old fathers reported by studies, few studies report no effects of paternal age on offspring quality (Heinze et al, 2018; Sparks et al, 2022). Others yet show that old fathers can produce larger (Aguilar et al, 2023; Mirrhosseini et al, 2014; Pappert et al, 2023), longer lived (Angell et al, 2022; Johnson et al, 2018; Krishna et al, 2012; Lee et al, 2019; Priest et al, 2002), and more fecund (Krishna et al, 2012; Sparks et al, 2022) offspring than young fathers (also see Kroeger et al, 2020 for beneficial maternal age effects). Some mechanisms have also been proposed to explain why old fathers produce higher quality offspring compared to young fathers. Males who mate at an old age are predicted to have, on average, longer lifespans than males who mate at younger ages (Mueller, 2004). However, this effect might be buffered in scenarios where old males are frail and die soon after mating (age-dependent frailty, Appendix 1), or when extrinsic mortality is high (Kokko and Lindstrom, 1996). However, if differences in survival between individuals are due to intrinsic reasons, older fathers would carry alleles that confer higher viability (Bowen et al, 2006; Chen and Maklakov, 2012; Kokko and Lindstrom, 1996; Reznick et al, 2004). Consequently, offspring born to older fathers would inherit these alleles, leading to old fathers producing longer-lived offspring (Beck et al, 2002; Brooks and Kemp, 2001; Hansen and Price, 1995; Kokko, 1998; Johnson and Gemmell, 2012; henceforth “viability selection hypothesis”).

The effects of paternal age at conception and paternal lifespan on paternal reproductive output and offspring lifespan are unlikely to be independent. Non-mutually exclusive processes (effects on male reproductive output: *reproductive senescence, selective disappearance, terminal investment*; effects on offspring quality: *Lansing effect, viability selection*), might co-occur and interact with each other to shape the life-history of fathers and their offspring. For example, higher paternal allocation toward producing offspring of high quality could come at the cost of lower allocation toward producing many offspring or fathers living longer (trade-offs hypothesis: e.g. Johnson et al, 2018; Travers et al, 2021). Different processes could further interact to shape patterns of ageing in animals. For instance, selective disappearance might lead to beneficial effects of advancing age on reproductive output until mid-life. However, from mid-to late-life, these age-dependent improvements could be outweighed (or balanced) by deleterious effects of senescence, leading to curvilinear shapes of ageing (e.g. Cooper et al, 2020; Jones et al, 2014; McCleery et al, 2008; Reid et al, 2010; Sanghvi et al, 2023; Torres et al, 2011), or no overall effects of age (e.g. Cooper et al, 2021). Only few studies (on wild populations) have attempted to disentangle the confounding effects of paternal age at conception and paternal lifespan (e.g. Reid et al, 2010; Torres et al, 2011). While valuable for their ecological relevance, these studies on wild populations have limited control over confounding factors such as maternal ageing and male mating history (a crucial modulator of male survival and reproductive output: Aich et al, 2022; Jones and Elgar, 2004; Partridge and Andrews, 1985; Paukku and Kotiaho, 2005).

Here, we use the fruit fly *Drosophila melanogaster* to manipulate paternal age at conception (PAC), and test its effects on paternal survival (aim 1), paternal reproductive output (aim 2), and the lifespans of sons (aim 3). We first investigated the influence of PAC on paternal lifespan and future survival (aim 1, Appendix 1). The relationship between PAC and paternal lifespan is assumed to be positive (H1A). However, this relationship might be flat or negative if males who mate when old are frail, thus incur more severe costs of mating and die soon after mating, than males who mate at a young age (H1B). Next, we investigated whether a male’s age at conception affects his reproductive output (aim 2, Appendix 2). In line with reproductive senescence (H2A) old males might have a lower reproductive output than young males. However, if male mortality is non-random and lifespan and reproductive output co-vary positively, males with low reproductive output might die earlier than males with high reproductive output (H2B), leading to an increase in reproductive output with advancing age (selective disappearance). Further, terminal investment could lead old males, who are close to dying, to invest more in a current reproductive event than young males (H2C). Finally, we investigate how PAC affects the lifespans of offspring, focussing on sons (aim 3, Appendix 2). Here, in line with Lansing effects (H3A), old fathers might produce shorter-lived sons than young fathers. However, if old fathers have alleles that confer higher lifespans, then sons born to old fathers might live longer than those born to young fathers (viability selection: H3B).

## Methods

### Stock population

To investigate how PAC affects paternal survival and reproductive output, as well as the lifespan of sons, we first set up a population of experimental males. To do this, we collected 300 virgin males (henceforth “unmated experimental males”) within 6 hours of eclosion on ice, and placed them in individual vials. These 300 males were collected from a stock population cage of lab-adapted, outbred, wildtype Dahomey *D. melanogaster* flies maintained in our lab since the 1970s. Unmated experimental flies were reared using a standard larval density method at 25°C and 45% r.h. (Clancy and Kennington, 2001). All flies in our experiment were maintained on a 12:12 hour light cycle, and fed with Lewis medium supplemented with ad libitum live yeast (Lewis, 1960). Under these conditions, male flies in our lab have median and maximum lifespans of ∼45 days and ∼90 days, respectively.

### Experimental design

We first generated different PAC treatments of experimental fathers. To do this, every two weeks (starting at 4 days of age), we chose between 25 to 37 surviving males from the 300 unmated experimental males (sample sizes in Table S1). In total, we generated a total of six PAC treatments, with males of the following ages (in days): 4, 18, 32, 46, 60, 74 (see Table S1 for sample sizes). We generated random numbers on MS Excel (‘randbetween’ function) to randomise our choice of males. The chosen experimental males were placed in a vial with a young (4 days old) virgin female of Dahomey background. Chosen experimental males were mated only once with the female, and males that did not mate within four hours (5 in total across all PAC treatments) were censored. Each mating was observed and mating latency and copulation duration recorded. After mating, the males were transferred into a new vial and kept individually until they died, and monitored daily to record their lifespans within 1 day of error. The mated females were left in the mating vial for another 24 hours following exposure to the male, to lay eggs, after which the females were discarded. These vials were left at 25°C and 45% r.h for 9 to 10 days, until offspring emerged (pupae in all vials emerged).

Within six hours of emergence, three male offspring (i.e. sons) were selected haphazardly from each parental pair and moved into a separate vial. All three sons from a parental pair were kept together until death. The remaining offspring from each parental pair were frozen at −20°C on the fourth day after eclosion began, and their numbers counted. We additionally recorded the survival (daily) of the unmated experimental males each day, to compare the survival of mated versus unmated males. All experimental flies, including unmated males and sons, were put in new food vials once every four days, using an aspirator.

### Data analysis

We used R v3.5.2 for all analyses (See HTML supplement for code and full model outputs), and packages *lme4* (Bates et al, 2014), *glmmTMB* (Brooks et al, 2014), *lavaan* (Rosseel, 2012), and *nlme* (Pinheiro et al, 2017) for building models. P values for all linear models were calculated using Satterthwaite’s method. All linear models were checked for model assumptions of normality and homoscedasticity of residuals using the *stats* package (R core team, 2012). All generalised linear models were checked for overdispersion using *DHARMa* (Hartig and Hartig, 2017). We interpreted main effects only when two-way interactions were non-significant (following Engqvist 2005), by fitting a model with the interaction removed. PAC was modelled as a continuous, not categorical variable, in all our models.

#### Aim 1: Effects of PAC on paternal survival

First, we investigated the effects of increasing PAC on paternal lifespan (see Appendix 1 and 2 for hypotheses and predictions). For this, we created a linear model with PAC as a fixed effect, and paternal lifespan as the dependent variable. Our data violated linear model assumptions of homogenous variance (i.e. lower variance in paternal lifespans for older PAC treatments). Thus, we specified a heterogeneous variance structure using the function *gls* (Pinheiro et al, 2017). A model with heterogenous variance structure was a better fit to the data than one without (ΔAIC = 7.817; ΔDF = 5).

Second, we tested whether mortality risk differed between mated (fathers) and unmated experimental males. For this, we compared survival probability between unmated experimental males and fathers who mated when young (4 days old). Young, rather than older fathers were chosen for this comparison because we wished to compare actuarial senescence patterns across the entire lifetime of mated versus unmated males. Using old fathers would not allow this comparison, because older fathers by definition, survived to older ages. We modelled data using Cox proportional hazards in the *coxme* (Therneau and Therneau, 2015) and *survival* (Therneau and Lumley, 2013) packages, with male lifespan as the dependent variable, and treatment (4 day old mated versus unmated) as a fixed effect.

Third, we compared whether mortality risk due to mating differed between young and old males, using a Cox proportional hazards model. For this, we used two approaches. In our first approach, we used a generalised linear model with binomial error distribution. Here, we compared the proportion of males that were alive (1) or dead (0) (dependent variable), in each PAC treatment (fixed effect), three days after mating. Here, we additionally included number of offspring each male produced as a fixed effect to test for potential trade-offs between paternal mortality risk and offspring production. However, differences in survival soon after mating, between different PAC treatments could, in principle, be explained by actuarial senescence alone. We thus used a second approach to specifically test whether mortality risk differed between old and young males due to mating *per se*, by removing effects of actuarial senescence. To do this, we compared survival probabilities between males who mated when 60 days (i.e. old PAC) old and males mated when 4 days old (i.e. young PAC), while only using data of males from both PAC treatments that survived beyond 60 days. Similarly, we also compared survival probabilities between males who mated when 74 days old versus 4 days old, but only using data on males from both PAC treatments that survived beyond 74 days.

#### Aim 2: Effects of PAC on paternal reproductive output

We tested the effect of PAC on paternal probability of siring an offspring (P_S_), and on the number of offspring sired (N_S_) when only considering fathers who sired offspring. To do this, we used a hurdle generalised linear model with truncated negative binomial error distribution (Brooks et al, 2017), and number of offspring as our dependent variable. A hurdle model allowed us to first, compare the P_S_ of fathers (i.e. zero inflation model). Then, using only data on fathers that sired offspring (conditional model), compare the N_S_ of fathers. PAC, paternal lifespan, their interaction, paternal copulation duration, and mating latency, were included as fixed effects. Copulation duration was included to account for males potentially producing more offspring due to transferring more sperm by copulating for longer (Dore et al, 2021). Latency was included to account for differences in female pre-mating preference for males, where females who take longer to mate (i.e. less prefer a male), might produce fewer offspring than females who take shorter to mate. In the same model, we also included a quadratic term for PAC, to test for curvilinear patterns of ageing. This model would allow us to test for the effects of reproductive senescence, as well as selective disappearance, by testing whether within each PAC treatment, fathers with longer lifespans sired more offspring than fathers with shorter lifespans. We conducted a sensitivity analysis to ensure that the patterns observed for reproductive ageing were not driven by the oldest age group (i.e. 74-day old males), by re-analysing the data but excluding the 74-day old males, as this age group had few data points (Table S1).

We then investigated whether observed patterns in paternal reproduction could be due to terminal investment, by testing whether old fathers who were closer to dying produced more offspring than old fathers who were less close to dying, or than young fathers. For this, we first calculated the number of days elapsed between a male’s death and the day he mated (henceforth called “days to death”). Then, using a generalised linear model with zero-inflated negative binomial error distribution, we tested how the interaction between days to death and PAC (fixed effects), affected the number of offspring produced (using all data, including fathers who produced zero offspring) by fathers (dependent variable). Here, we additionally included paternal copulation duration as a fixed effect.

#### Aim 3: Effects of PAC on lifespans of sons

We investigated the effects of PAC on the lifespans of sons, and whether the observed effects were due to fathers in older PAC treatments living longer (i.e. viability selection). For this, we used three approaches. In our first approach, we tested how PAC affected the lifespans of sons (dependent variable) using a linear model. Here, we included PAC as fixed effect and paternal ID as a random effect. We also included the number of offspring produced as fixed effect to test for trade-offs between paternal investment in offspring lifespan versus number (Johnson et al, 2018). Our models had heterogenous variance structure specified, as this structure yielded a better fit to the data than a model without heterogenous variance (ΔAIC= 25.66; ΔDF= 1).

In our second approach, we tested whether PAC had a significant effect on the lifespans of sons, after accounting for the variance explained by paternal lifespan. We used this approach to test whether the effect of PAC on lifespans of sons, observed in our first approach, was mediated by paternal lifespan (i.e. indirectly) as predicted by the viability selection hypothesis. We first built a model with paternal lifespan as a fixed effect, paternal ID as a random effect, and the lifespan of each son as our dependent variable. Then, we used residuals from this model as the dependent variable, and included PAC and the number of offspring produced by fathers as fixed effects, and paternal ID as a random effect. This model had a heterogeneous variance structure specified as described above.

In our third approach, we conducted a path analysis using a structural equation model in the package *lavaan* (Rosseel, 2012). This approach was used to obtain a better understanding of the path of causality for how PAC affects lifespans of sons. We modelled the direct effect of PAC on son lifespan and its indirect effect via the influence of paternal lifespan. In this model, we also included covariances between number of offspring produced by fathers with paternal lifespan and with lifespans of sons, to account for trade-offs between offspring quality and number, and between investment in reproduction versus survival by fathers.

## Results

### Aim 1: Effects of PAC on paternal survival

Increased paternal age at conception (PAC) was associated with an increase in the average lifespan of fathers (t= 3.072; P= 0.003, R^2^ = 4.3%, Figure 1). Paternal lifespan increased by 0.125 days with an increase of one day in PAC.

**Figure 1:**
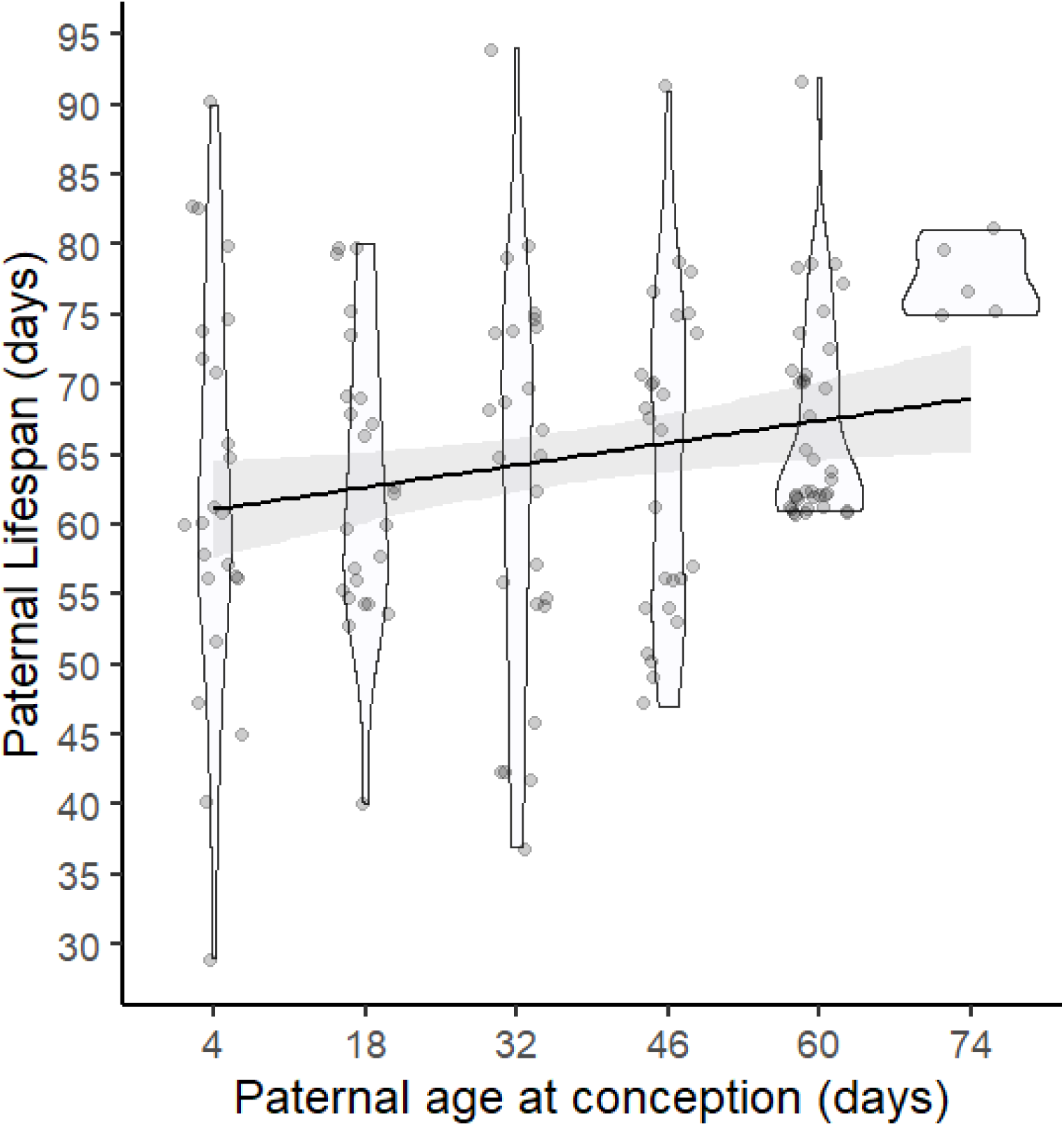
Males who conceive at older ages have longer lifespans on average. Violin plots show smoothed distribution of data, while dark line and shaded areas show regression line and 95% C.I. Lower Y axis limit set to 25 because no deaths occurred before 25 days of age.

There was no significant difference in survival probabilities between the unmated experimental males, and those who mated at 4 days old (P = 0.12, Figure 2 and 3A). When comparing only males who mated, males in older PAC treatments had a lower probability of surviving beyond three days after mating, than males in younger PAC treatments (z = -3.544, P < 0.001, Table S2, Figure S1). For instance, within 3 days after mating, males who mated when ‘old’ (i.e. at 60 or 74 days old) experienced >50% mortality, compared to no deaths in ‘young’ PAC treatments (i.e. 4, 18, and 32 days old). While this result could be due to age-specific effects of mating stress, it could also be a consequence of actuarial senescence (older PAC had an overall lower age-dependent survival probability than young PAC treatments: z = -2.067, P = 0.038, Figure S2). We thus further tested whether the difference in mortality rates between old and young PAC treatments was specifically due to mating. Males who mated at 60 days of age had a significantly lower probability of surviving past 60 days of age than males who mated at 4 days of age, when only data on males that survived past 60 days of age in both PAC treatments were used (P = 0.045, Figure 3A). Similarly, males who mated at 74 days of age were less likely to survive past 74 days of age than males who mated at 4 days of age (P = 0.066, Figure 3B). Collectively, these results indicate that while advancing PAC is associated with fathers having longer lifespans, the strength of this effect is buffered by old fathers experiencing higher mortality associated with mating stress, compared to fathers who mated when young.

**Figure 2:**
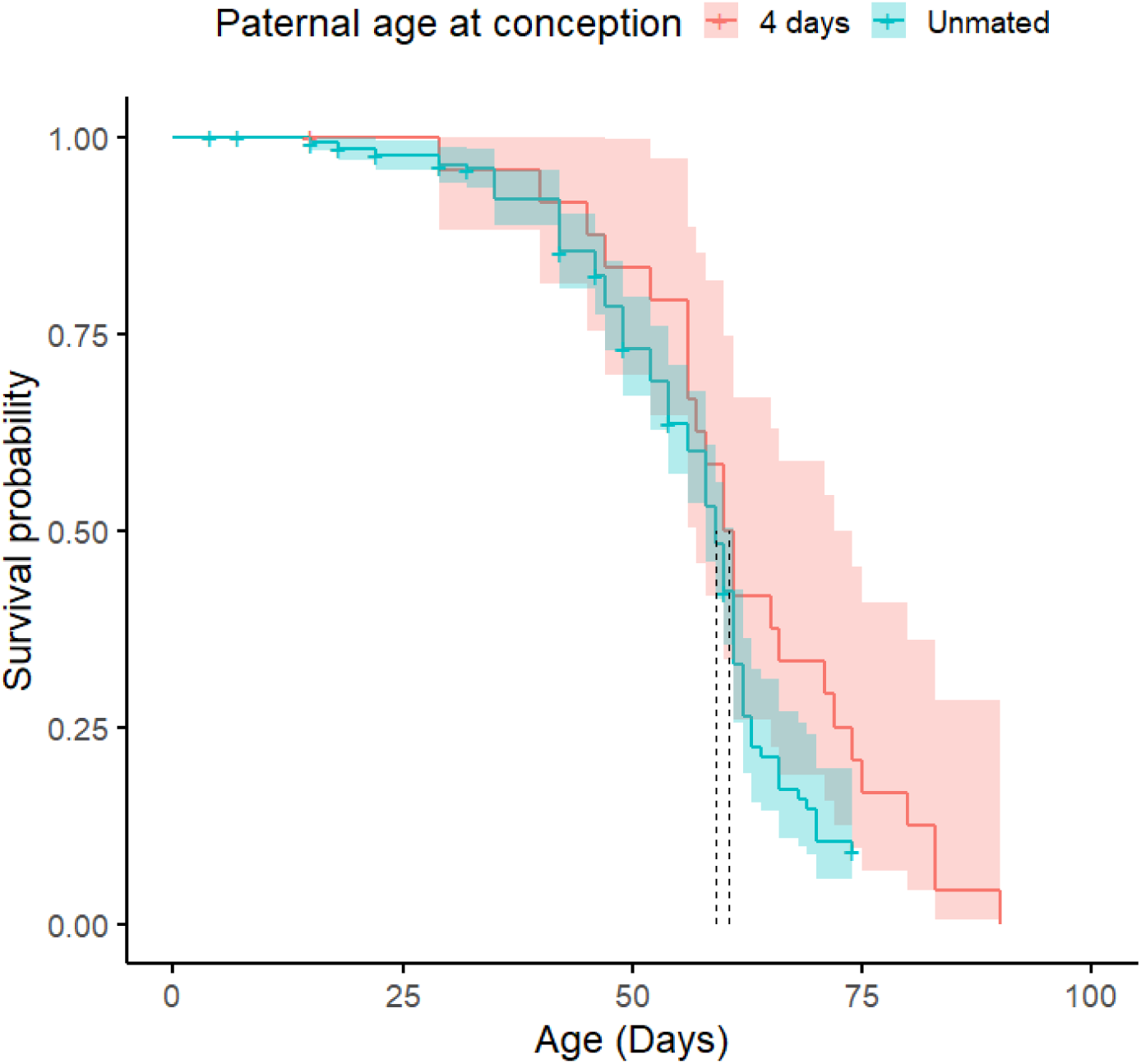
Males who mated at 4 days of age did not have a significantly different survival probability compared to the unmated experimental males. “+” indicates censored individuals. Shaded area shows 95% C.I. Dotted lines show age at median survival probability.

**Figure 3:**
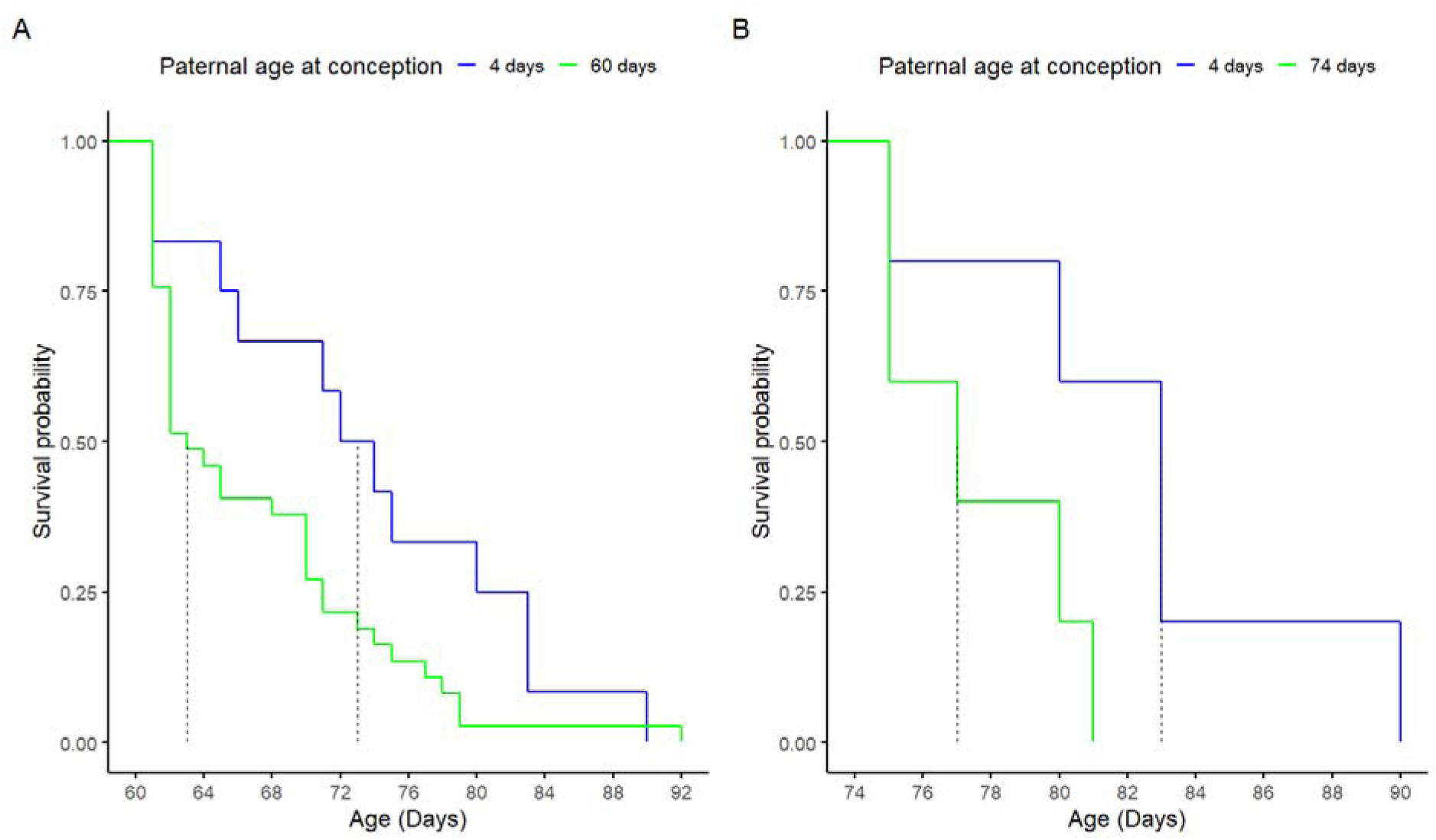
A-Males who mated at 60 days of age had a lower survival probability than males who mated at 4 days of age, using only data on males that survived past 60 days of age for both groups. B-Males who mated at 74 days of age had a lower survival probability than males who mated at 4 days of age, using only data on males that survived past 74 days of age for both groups. Dotted lines show age at median survival probability.

### Aim 2: Effects of PAC on paternal reproductive output

We found a significant interaction between PAC and paternal lifespan, on the probability of siring offspring (i.e. P_S_ : z = -2.336, P = 0.020, Table S3). Specifically, P_S_ increased with paternal lifespan, but only for fathers who conceived at older ages (Figure 4A). When investigating variation in N_S_ (i.e. number of offspring sired by fathers, who produced at least one offspring), we found no significant effects of the interaction between PAC and paternal lifespan (z = 0.562, P= 0.574, Table S3), and no significant effect of paternal lifespan (z = 0.963; P = 0.336) on paternal N_S_. However, we found a significant quadratic effect of PAC on N_S_ (z = -3.902, P < 0.001, Figure 4B). Our sensitivity analysis (i.e. a model without data from 74 day old PAC) also showed a significant quadratic effect of PAC on N_S_ (z = -2.084, P = 0.037). However, the shape of PAC on N_S_ from the sensitivity analysis was shallower (Figure S3, S4) than the original model that included data from 74 day old males. This suggests that a decline in N_S_ only becomes apparent in fathers who mate after 60 days of age.

**Figure 4:**
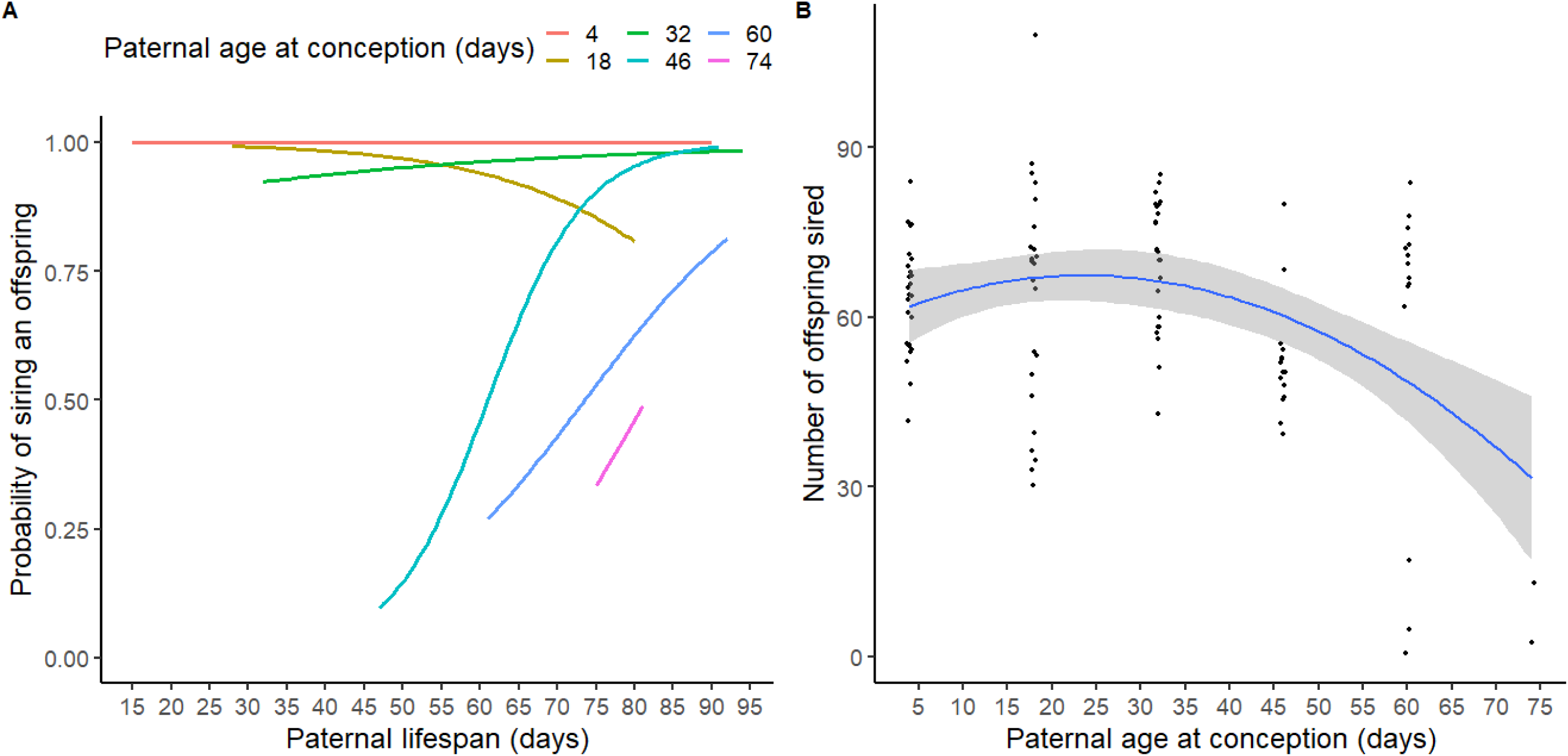
A: Paternal age at conception interacted with paternal lifespan to affect the probability of siring an offspring (P). B: Paternal age at conception affected the number of offspring sired (N : after excluding fathers who did not produce offspring) in a quadratic way. Shaded areas represent 95% C.I.

We found a significant interaction between days to death (i.e. number of days elapsed between mating and death) and PAC to affect number of offspring sired by fathers. Specifically, old males who died soon after mating sired fewer offspring than old males who died later after mating, or than young males (z = 2.201, P = 0.027; Table S4). Overall, days elapsed between death and mating did not significantly influence the number of offspring a male sired (z = 0.167, P = 0.867). Collectively, these results indicate that reproductive senescence in fathers becomes apparent only in late-adult life; that life-history traits of survival and reproductive output show pleiotropic interactions that depend on the age of the father; and that there is no evidence for terminal investment or selective disappearance.

### Aim 3: Effects of PAC on lifespans of sons

In our first approach, we tested for the effects of PAC on lifespans of sons without accounting for effects of paternal lifespan. We found a significant positive effect of PAC on the lifespans of sons (t = 2.532; P= 0.013; Table S5, Figure 5, Figure S5). Lifespans of sons increased by ∼0.1 days with each day of increase in PAC. The number of offspring produced by fathers had no effect on the lifespans of sons (t = 0.054, P = 0.957). In our second approach, we tested for the effects of PAC on residuals from a model that first tested for effects of paternal lifespans on lifespans of sons. Here, we found no significant effects of PAC on these residuals (t = 1.532; P = 0.129). In our third approach (Figure S6; Table S6), our path analysis suggested significant direct effects of PAC on sons’ lifespan (z = 2.443, P = 0.015) and on paternal lifespan (z = 4.662, P < 0.001). This path analysis also revealed marginally non-significant effects of paternal lifespan on sons’ lifespan (z = 1.920, P = 0.055), as well as marginally non-significant indirect effects of PAC on sons’ lifespans via the effect of paternal lifespan (z = 1.776, P = 0.076). Collectively, our results for aim 3 indicate that the positive effects of advancing PAC on the lifespans of sons is partly due to older fathers living longer. This result is in line with the viability selection hypothesis rather than Lansing effects, and is not due to trade-offs between paternal reproductive output and sons’ lifespans.

**Figure 5:**
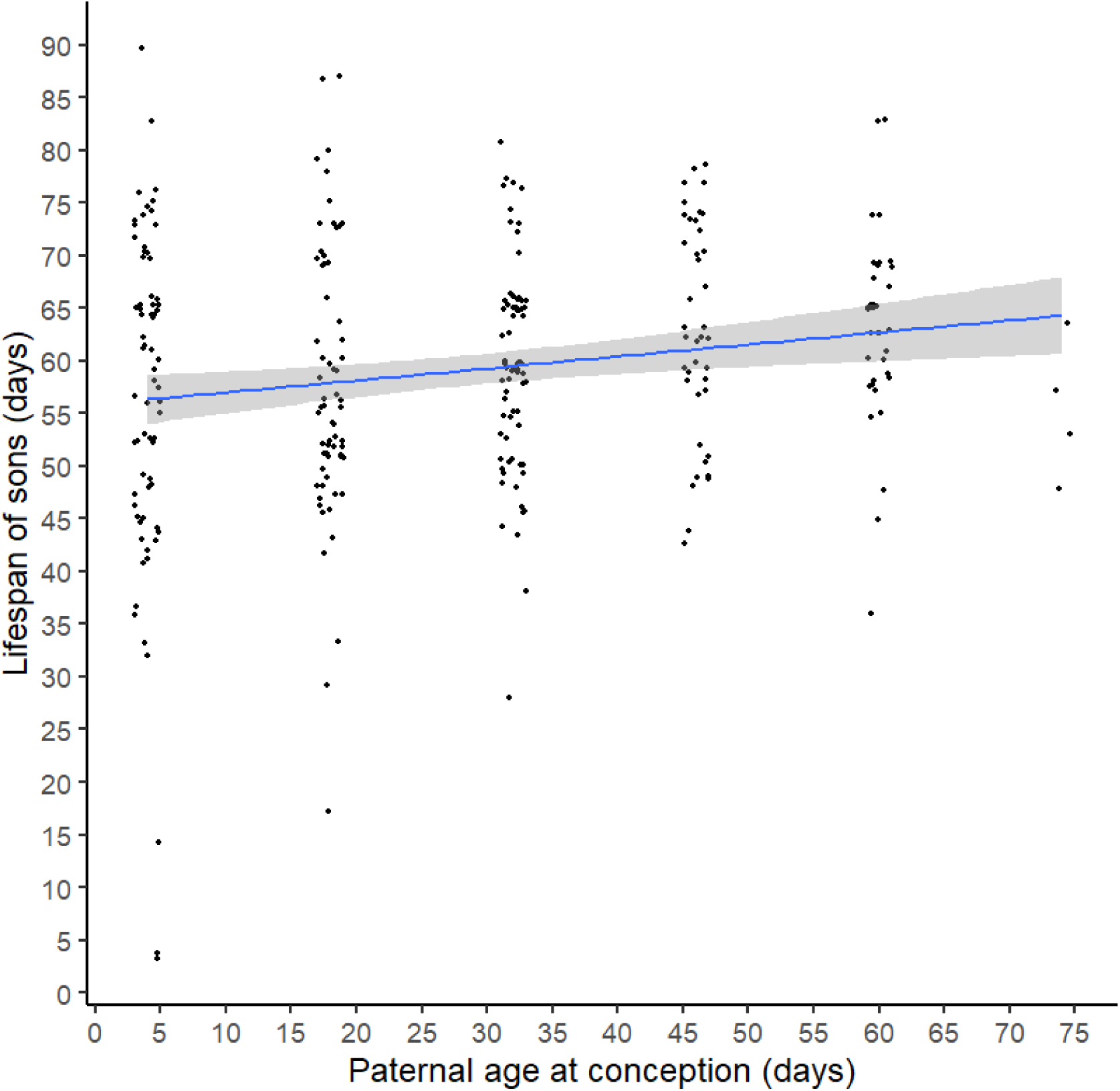
Increase in paternal age at conception led to an increase in the lifespans of sons. Each dot represents the lifespan of a single son, shaded areas show 95% C.I. See figure S5 for means and SE.

## Discussion

Paternal age at conception can affect paternal survival and reproductive success, and the fitness of offspring in complex ways, with several hypothesized outcomes. Studying the effects of paternal age therefore requires simultaneously testing for multiple, non-mutually exclusive processes (e.g. reproductive senescence, selective disappearance, terminal investment, Lansing effect, viability selection, trade-offs). Currently, there is a scarcity of studies adopting such an approach. We used experiments in fruit flies to investigate how paternal age at conception (PAC) affects the survival (aim 1) and reproductive output (aim 2) of fathers, and the lifespans of their sons (aim 3), and show that the effects of PAC are more complex than predicted by any single hypothesis (See Appendix 1 and 2 for predictions for each hypothesis).

### Aim 1: Effects of PAC on paternal survival

We found that while increased PAC was associated with higher paternal lifespans, mating itself was related to increased mortality in older fathers. Mating is known to be energetically and physiologically costly to individuals, and can often reduce overall longevity (Heinze, 2016; Kemp and Rutowski, 2004; Koppik et al, 2018; Partridge and Andrews, 1985; Paukku and Kotiaho, 2005; Perry and Tse, 2013; Sanghvi et al, 2021; Scharf et al, 2013). We found no differences in the overall survival of unmated males and that of the fathers who mated when young. However, we found that fathers who mated at older ages had higher mortality than fathers who mated when younger (see Clinton and Le Boeuf, 1993, who show the opposite in seals), even after removing the effects of actuarial senescence, which has not been previously reported. This result is likely due to old males being frailer and physiologically more vulnerable than young males (e.g. Bissett and Howlett, 2019; Minois and LeBourg, 1999), such that the cost of mating affects the survival of old males more than young males.

### Aim 2: Effects of PAC on paternal reproductive output

We found significant interactive effects of paternal lifespan and PAC on paternal P (the probability of siring an offspring). Specifically, paternal lifespan covaried positively with paternal P, but only for fathers who conceived at older ages. This result could be because most (>92%) males in the three youngest PAC treatments sired offspring, and infertility became apparent only in older PAC treatments. This higher variance in P within older PAC treatments could have led to a greater opportunity for covariance between lifespan and P. While the covariance between lifespans and P could have also been negative, a positive covariance would be expected if reproductive output is condition-dependent (Bonduriansky and Brassil, 2005; Chen et al, 2016; Sultanova et al, 2020). Thus, for an older male, ceasing of offspring production could be a reliable indication of the individual’s proximity to death. However, for young males this would not be so, because of low variance in P. Future studies could investigate whether these patterns hold true, in systems where variance in reproductive success is the same for old and young males.

We obtained no evidence for the terminal investment hypothesis. Specifically, older males who died soon after mating produced fewer offspring than older males who died later, or than young males. This result is inconsistent with some studies in birds and insects, that have found older males (Duffield et al, 2018; Farchmin et al, 2020; Velando et al, 2006) and females (Creighton et al, 2009; Froy et al, 2013; Part et al, 1992) to invest terminally in reproduction. However, males in these studies were also immune-challenged, which can cause individuals to perceive themselves as being diseased, which was not the case in our study. We additionally found no evidence for an overall effect of paternal lifespan on the number of offspring sired by a male (N), and only found an effect of paternal lifespan on P for older PAC groups. Evidence for selective disappearance would require detecting positive covariances between paternal lifespan and N or P across all PAC groups. Our results thus suggest that selective disappearance cannot explain these patterns.

Our results showed some evidence for reproductive senescence (also seen in other fruit fly studies: Grotewiel et al, 2005; Ruhmann et al, 2018; Sepil et al, 2020). Reproductive senescence in our study was driven by old males being less likely to sire an offspring, and the oldest males (i.e. 74 days old) siring the fewest numbers of offspring. Overall, our results suggest that male reproductive output increases from mid-to adult-life, and reproductive senescence onsets in late-adult life (Baudisch and Stott, 2019; Jones et al, 2014; Sanghvi et al, 2023). Future studies could investigate the extent to which the patterns of reproductive senescence observed in our study are due to declines in sperm number (Sepil et al, 2020; Gasparini et al, 2010, 2019), sperm performance (Dean et al, 2010; Vuarin et al, 2019), seminal fluid quantity (Reinhardt et al, 2019), or changes in seminal fluid composition (Fricke et al, 2023).

### Aim 3: Effects of PAC on lifespans of sons

Older fathers produced longer-lived sons, a result that is not in line with the Lansing effect (Crow, 2003; Lansing, 1947; Monaghan et al, 2020; Sharma et al, 2015; Wylde et al, 2019). Our results contrast with other studies in fruit flies that show a Lansing effect (e.g. Mossman et al, 2019; Price and Hanson, 1998), whereby offspring born to older fathers have lower survival. There could be several reasons for this discrepancy.

First, fathers in our experiment were kept individually isolated, without exposure to rival males, whereas Mossman et al (2019) and Price and Hanson (1998) kept males in single-sexed groups. Exposure to rivals has been shown to cause *Drosophila* males to invest more in sperm production (Bjork et al, 2007; Hopkins et al, 2019) at the cost of reduced investment in maintaining sperm quality (Koppik et al, 2023; Silva et al, 2019) compared to males kept individually. Thus, it is possible that fathers in our experiment would have been able to prevent deleterious paternal age effects mediated via deterioration in sperm quality, due to being kept individually. Second, lack of exposure to rival males in our study could have reduced the rate of sperm production, thus germline cell division and mutation rates in fathers (de Manuel et al, 2022; Crow, 2000; Girard et al, 2016; Monaghan and Metcalfe, 2019), compared to studies showing Lansing effects that kept males in groups. This possible reduction in germline mutation rate could further buffer the effects of paternal age. Third, sons in our experiment were not mated. Life-history theory predicts trade-offs between allocating energy to somatic maintenance versus reproduction (Lemaitre et al, 2015; Stearns, 1989). Being kept unmated could have allowed sons to invest in somatic maintenance, possibly masking deleterious paternal age effects, and future studies can test this by repeating our experiments but having another treatment where sons mate. Fourth, we sampled fathers at extremely old lifespans (i.e. up to 74 days, representing >75% of maximum lifespan). However, studies that have shown deleterious paternal age effects in *D. melanogaster* sampled ‘old’ fathers between 14 and 45 days of age (e.g. Mossman et al, 2019; Nystrand and Dowling, 2014; Price and Hanson, 1998; Sepil et al, 2020). It is possible that at ages as extreme as in our study, deleterious effects of PAC are outweighed by positive effects of selection on paternal viability.

Our results provided some support for the viability selection hypothesis (Brooks and Kemp, 2001; Hansen and Price, 1995; Kokko, 1998; Johnson and Gemmell, 2012). Specifically, fathers that mated when older had longer lifespans, and produced sons with longer lifespans. When effects of paternal lifespans on sons’ lifespans were removed (i.e. second statistical approach), PAC no longer had a significant effect on the lifespans of sons. These results suggest that significant positive effects of PAC on lifespans of sons were driven by paternal lifespans. There could however, be other mechanisms which could explain the direct effect of PAC on the lifespans of sons. For instance, poor quality offspring of old fathers, could have experienced death at the larval or pupal stage, thus eclosed sons representing a biased sample of high-quality offspring (e.g. Sanghvi et al, 2022). Unlike Johnson et al (2018), we found no evidence that increased lifespans of sons from old males was due to trade-offs between paternal investment in reproductive output versus investment in offspring quality (i.e. increased lifespan of sons). Our results overall suggest that increasing PAC selects for fathers that have alleles which confer higher viability, which are then inherited by the sons of older fathers. Some previous studies have found positive effects of PAC on offspring lifespans (Angell et al, 2022; Johnson et al, 2018; Krishna et al, 2012; Lee et al, 2019; Priest et al, 2002). However, to our knowledge, ours is the first to formally test the viability selection hypothesis, by investigating separately, the direct as well as indirect effects (via paternal lifespans) of PAC, on offspring lifespans. Future studies could measure reproductive output of sons, and fitness-components of daughters, because paternal age can affect sons versus daughters differently (e.g. Angell et al, 2022; Bouwhuis et al, 2015; Priest et al, 2022; Schroeder et al, 2015; Sparks et al, 2022). Collectively, our results indicate that being born to old fathers need not be deleterious to offspring lifespan.

## Conclusions

Our study simultaneously tests various mechanisms that may link PAC to paternal survival (age-dependent frailty) and reproductive output (reproductive senescence, selective disappearance, terminal investment), and offspring lifespans (Lansing effect, viability selection, trade-offs between paternal investment in offspring quality versus quantity). We show that mating at older ages can lead to an increased risk of mortality for fathers, thus buffering the magnitude of positive covariances between PAC and paternal lifespans. We further reveal that positive pleiotropy between survival and reproductive output depends on the age of the father. We also find that reproductive ageing patterns of males is quadratic in shape, with declines in reproductive output occurring in late-life. Lastly, we show that advancing PAC has a positive effect on the lifespans of sons, and that this effect is likely due to older fathers having longer lifespans. We recommend that future studies employ a broader theoretical framework that weighs costs of ageing against its direct and indirect benefits, to better understand organismal health and life-history. Fitness benefits of old fathers producing longer lived sons might offset the fitness costs of reduced fertility in old fathers. If so, this offset could lead to the evolution of female preference for old males (Beck and Powell, 2000; Beck and Promislow, 2007; Beck et al, 2002; Johnson and Gemmell, 2012; Kokko and Lindstrom, 1996), and future studies can investigate this exciting but relatively unexplored avenue of research.

## Acknowledgements

We thank Devi Satarkar and Jinlin Chen for checking survival of flies over Christmas. We are also grateful to Chris Terry, Sam Gascoigne, Ellie Bath, and Sumali Bajaj for statistical advise, as well as Alex Kacelnik and members of the fly lab for helpful discussions.

## Data archiving

Data and all associated code are available on OSF with DOI: 10.17605/OSF.IO/38CXN at https://osf.io/38cxn/. Supplementary information is provided along with this manuscript.

## Supplementary information

## Appendix 1: Effects of paternal age on paternal survival (hypotheses and predictions)

It is commonly assumed that as age at conception increases, lifespan should increase, i.e., fathers who mate at an older age should have higher lifespans (H1A). A corollary prediction is that at older age groups, there should be lower variances in lifespan. These predictions are based on the fact that individuals who mate at an older age have a lower limit for what lifespans they can have, because they have by definition survived to the age at which they conceive. For instance, if males mate at age 60 days, all males have survived until at least 60 days of age. If all individuals have a similar upper limit to lifespans, irrespective of the age at which they mate, then older ages of conception would have lower variance but higher means in lifespans. This leads to the effects of age at conception on lifespan to look as follows (Figure A):

**Figure A:**
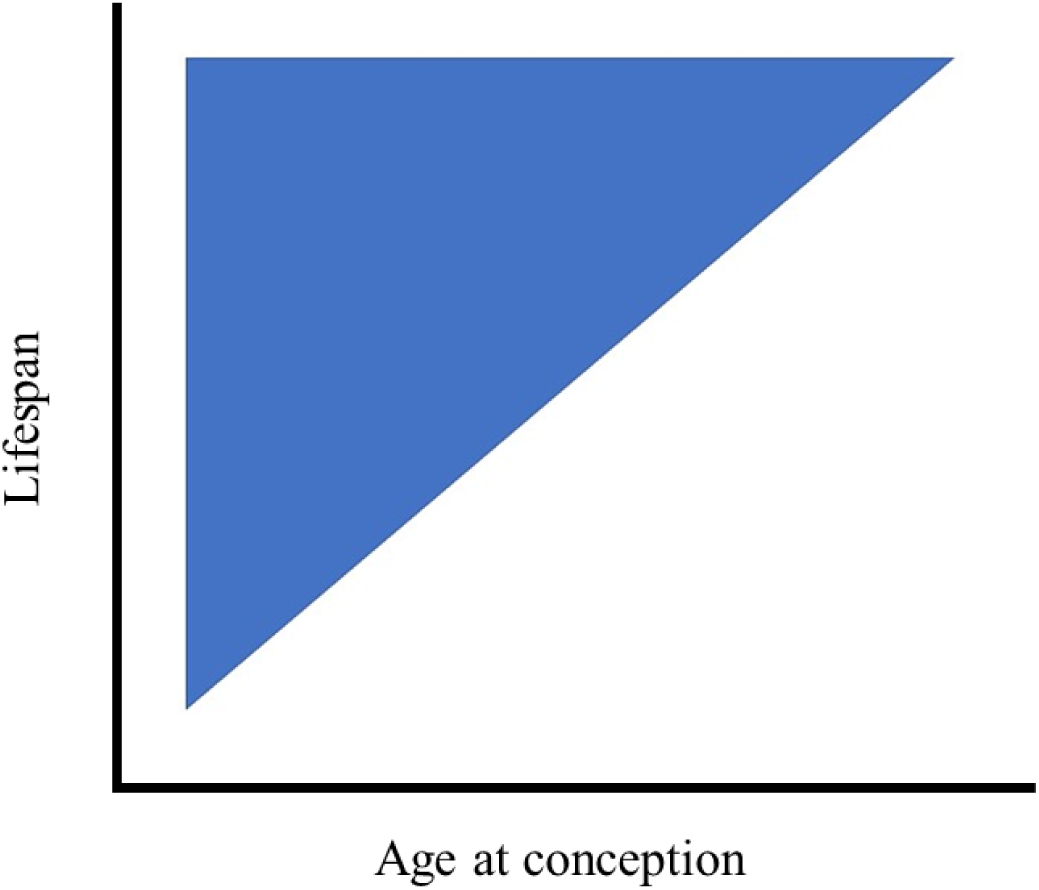
Advancing age at conception leads to higher means but lower variance in lifespan of older individuals

However, this prediction (corresponding to H1A) has two assumptions which are not usually explicitly tested. The first assumption is that mortality onsets from birth, thus all age classes at which individuals conceive offspring, would have experienced some mortality. However, it is possible that mortality in a population does not onset until a certain age (e.g. if survival curves are sigmoid shaped), thus, the lower limit for lifespans for various ages at conception might be the same. For instance, if in a population, males mate at ages 10, 20, 30, 40 …. to 90 days of age, but no males die until 40 days of age, then for males who mate at ages 10, 20, 30, and 40 days, the bottom limit of lifespan will be 40 days of age. This can lead to effects of age at conception on lifespan to look as follows (Figure B):

**Figure B:**
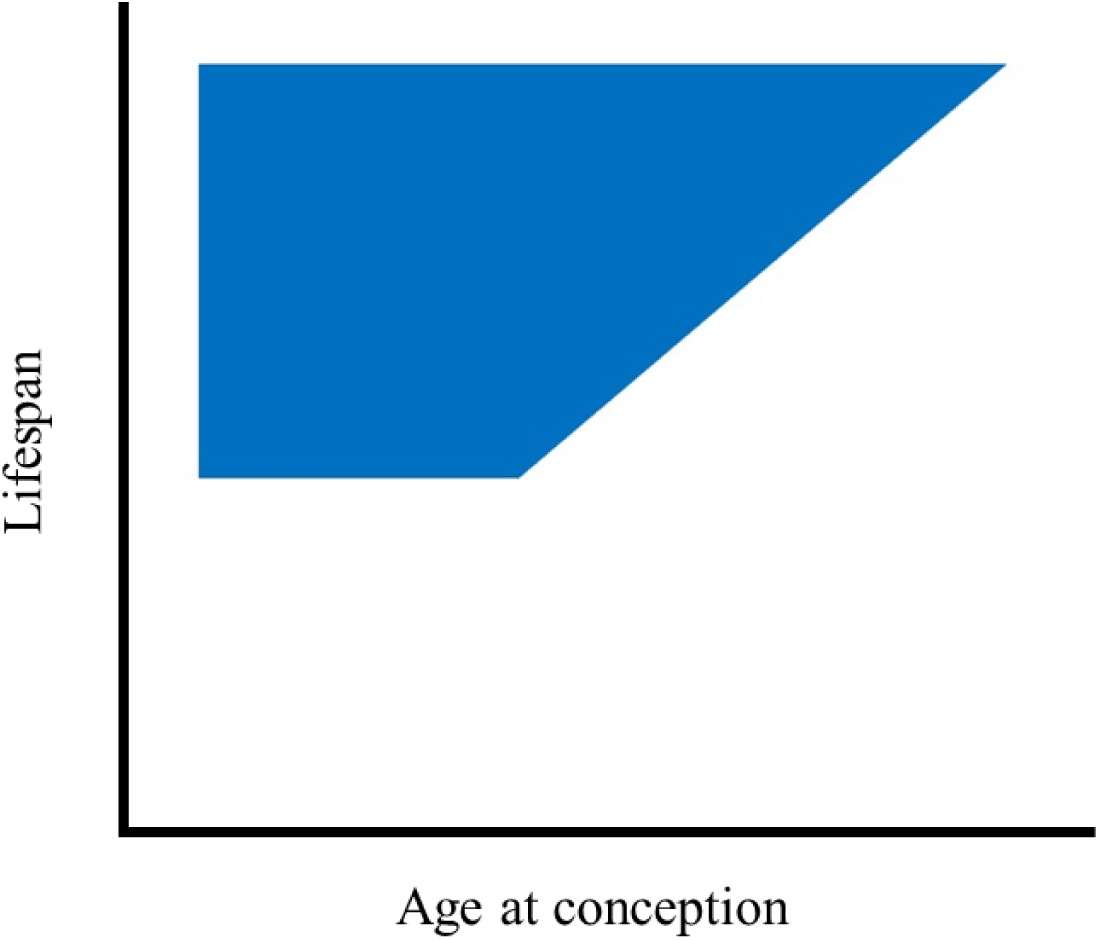
Advancing age at conception leads to higher means but lower variance in lifespan of older individuals, however, mortality does not occur until a certain age, thus age at conception groups until that age have the same average lifespans.

The second assumption is that the upper limit for lifespans across all ages of conception is the same. That is, irrespective of at what age individuals reproduce, they have the same probability of surviving until the upper limit of lifespan for that population. This however might not always be true (H1B). For instance, if older males are frailer than younger males, males mated when old might have a lower ceiling for longevity. An example of this is if mating related stress increases mortality in males who are more vulnerable (e.g. old mated males), thus reduces lifespans of males who mate when old but not when they mate young. This can lead to effects of age at conception on lifespan to look like Figure C.

**Figure C:**
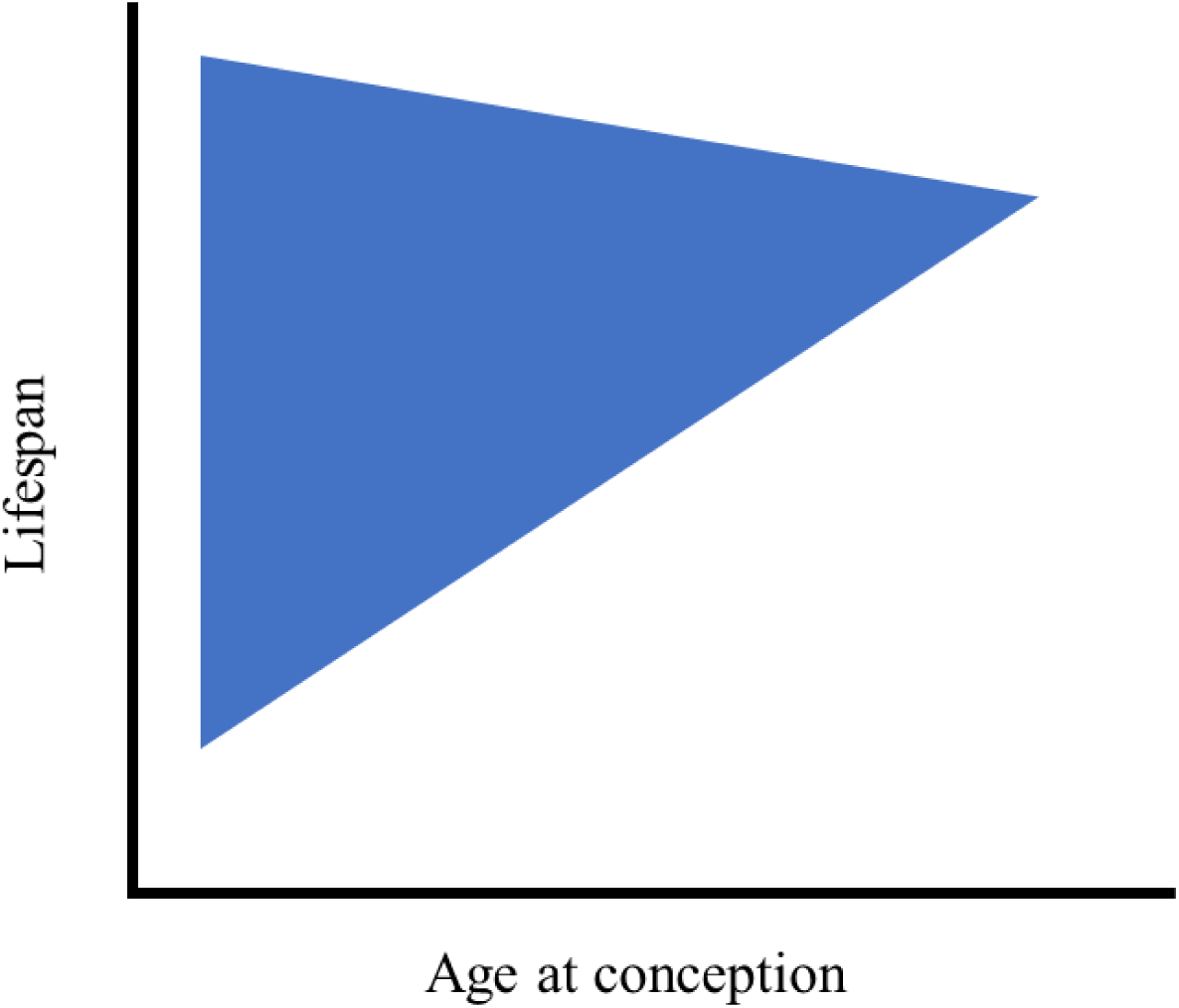
Advancing age at conception leads to higher means but lower variance in lifespan of older individuals. However, males who mate when older experience higher mating-related stress due to being frailer.

Combining the effects of assumption 1 and 2, we get a complex relationship between age at conception and lifespan (Figure D), which is quite different from the commonly held prediction in Figure A.

**Figure D:**
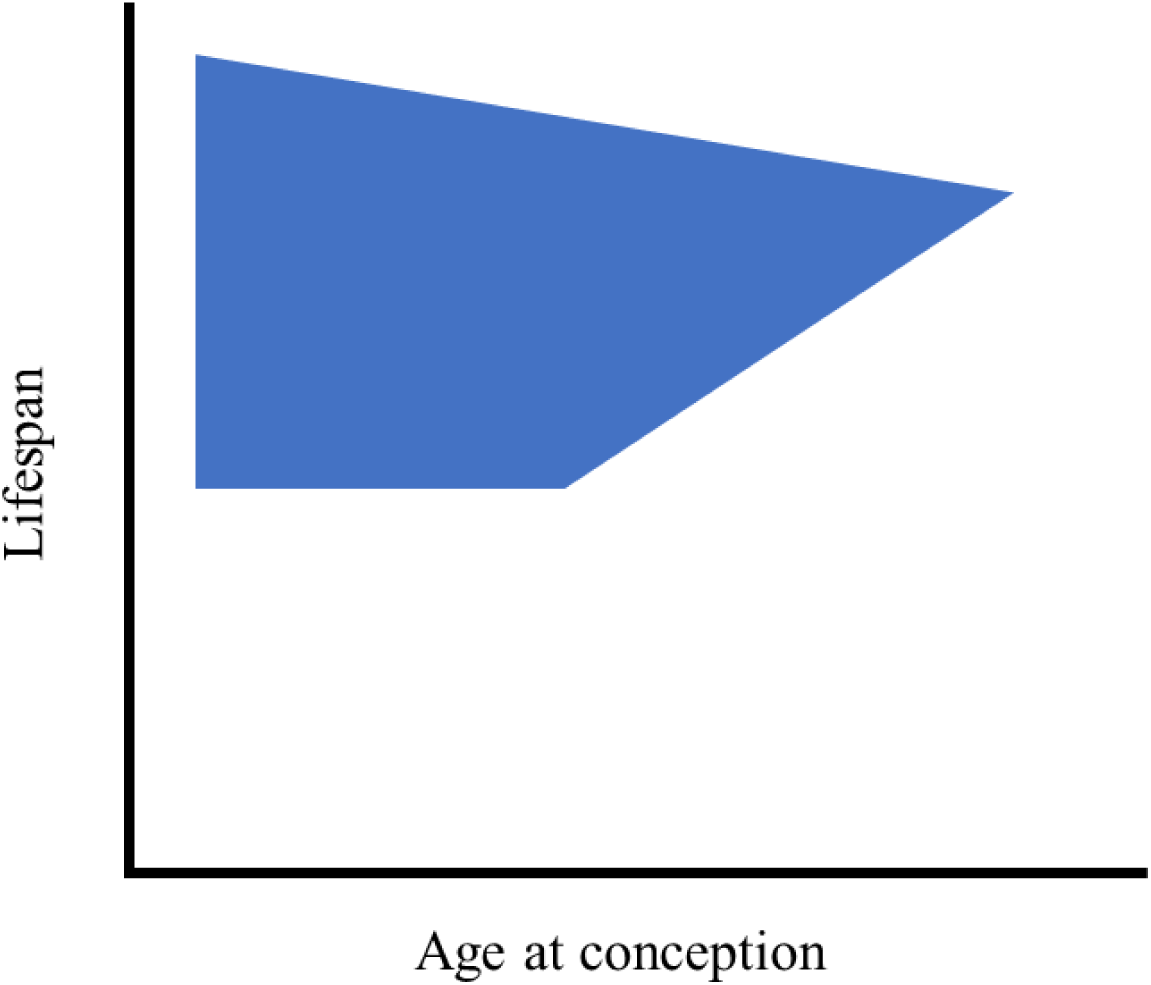
Advancing age at conception leads to higher means but lower variance in lifespan of older individuals. However, males who mate when older experience higher mating-related stress due to being frailer, and mortality does not occur until a certain age, thus age at conception groups until that age have the same average lifespans.

## Appendix 2: Graphical predictions for different hypotheses in each of the three aims, for how paternal lifespan, PAC, paternal reproductive output, and offspring lifespans, might be linked

### Note that presented graphs are hypothetical

1. Aim 1: Effects of PAC on paternal survival (also see Appendix 1)

a. Hypothesis 1: Age-dependent frailty to mating Prediction: If mating does not affect survival of males, then unmated males, and males who mate when young or when old, should all have same lifespans.

**Figure E:**
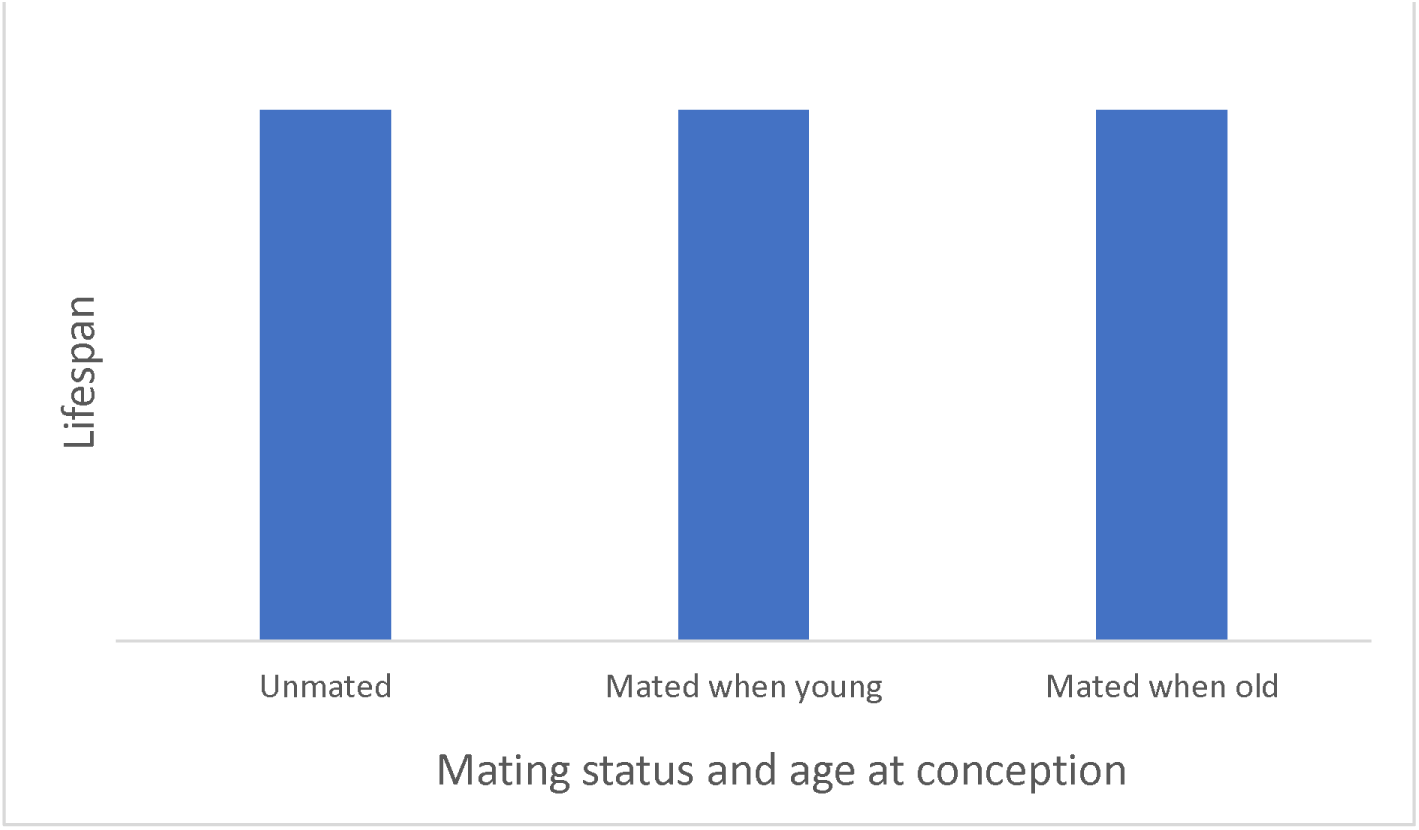
No effects of age and/or mating status to affect lifespan.

If mating-stress reduces the survival of males irrespective of the age at which males mate, then unmated males should have higher lifespans than mated males.

**Figure F:**
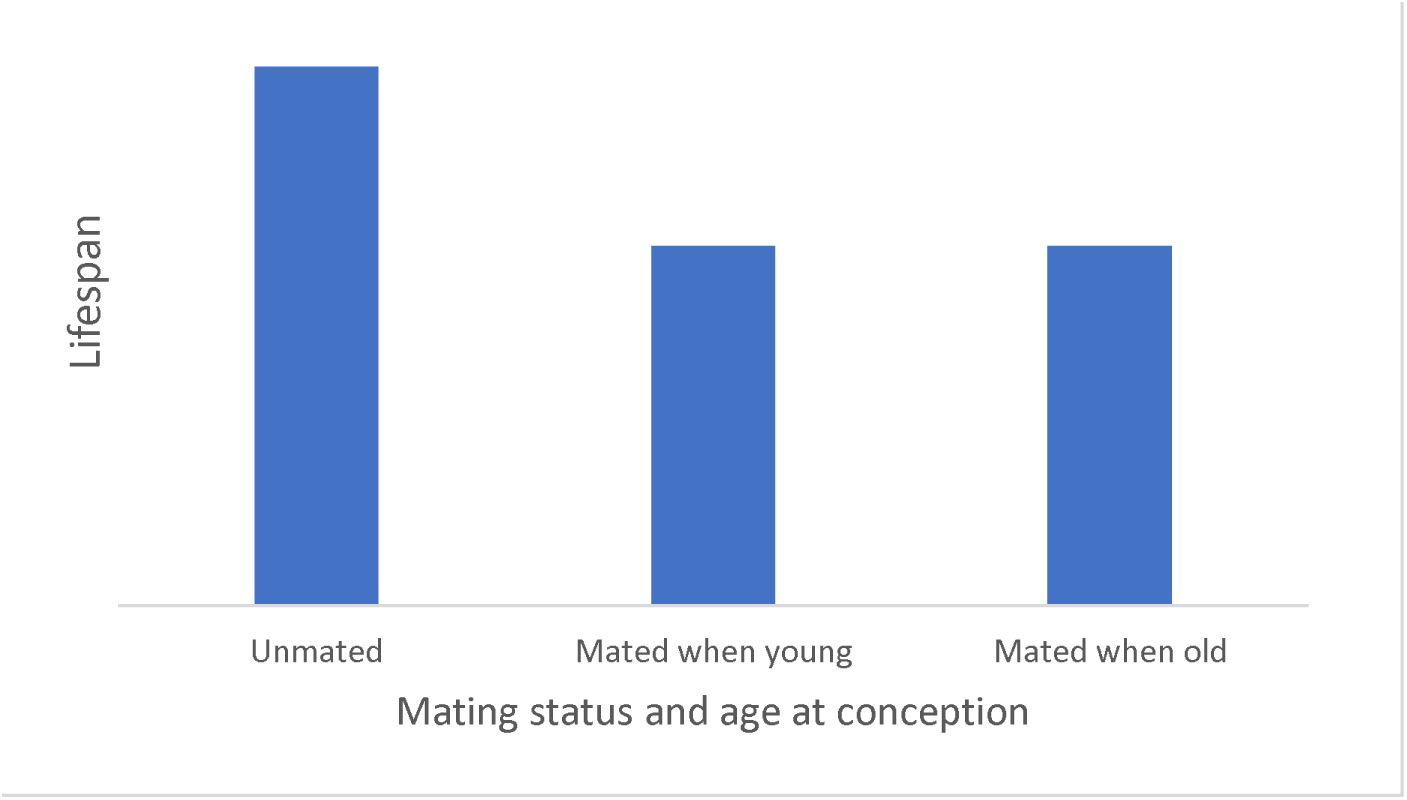
Mating status, but not age, affects lifespan, with unmated males living longer.

If old males are frailer and more vulnerable to mating stress, then mating-related stress should reduce the survival of males who mate when old, but not when young.

**Figure G:**
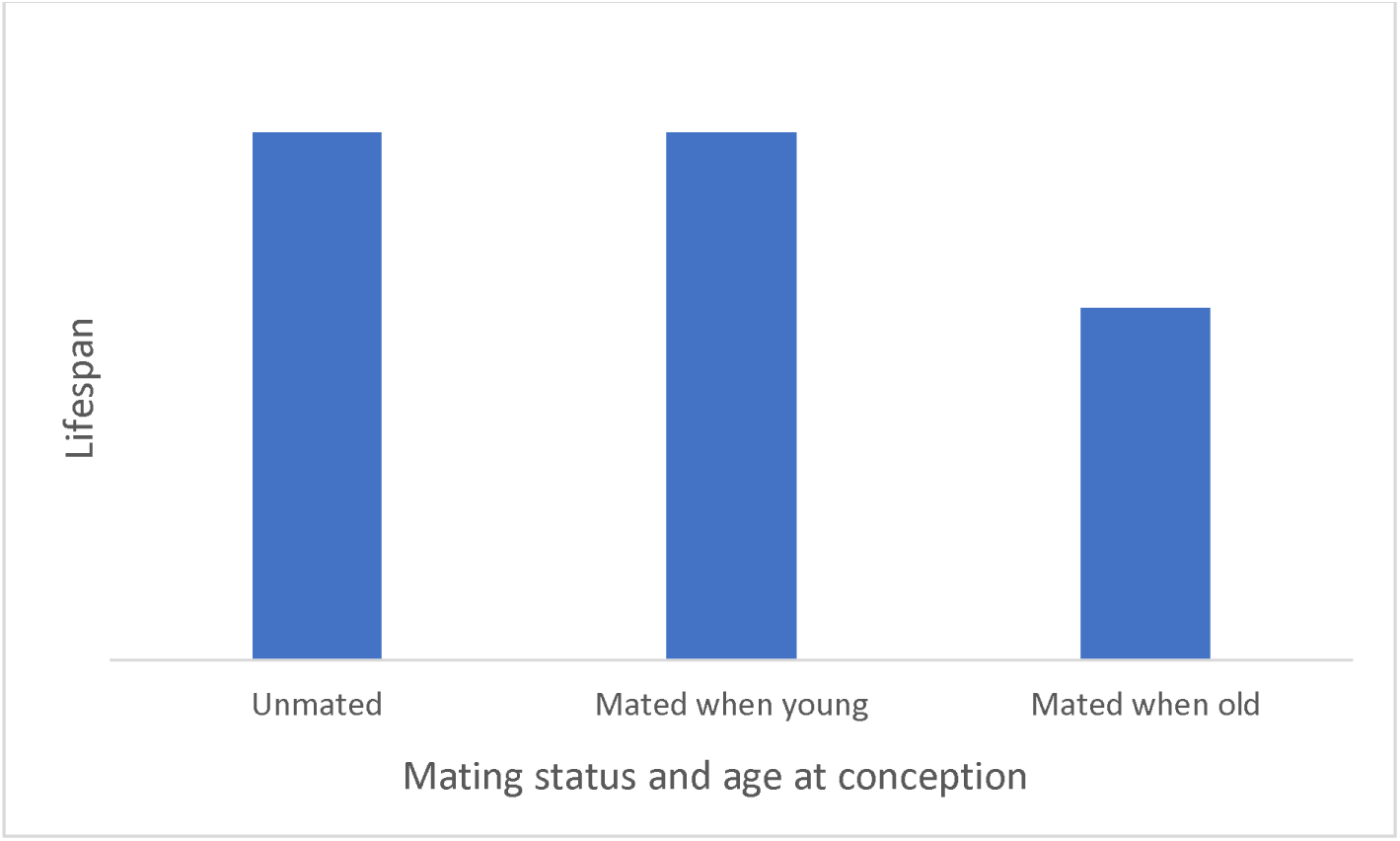
Mating status interacts with age to affect lifespan, with mated, older males living the shortest.

2. Aim 2: Effects of PAC on paternal lifespan

a. Hypothesis 2A: Reproductive senescence Prediction: If reproductive senescence occurs, old males should have lower reproductive output than young males. **Figure H:**
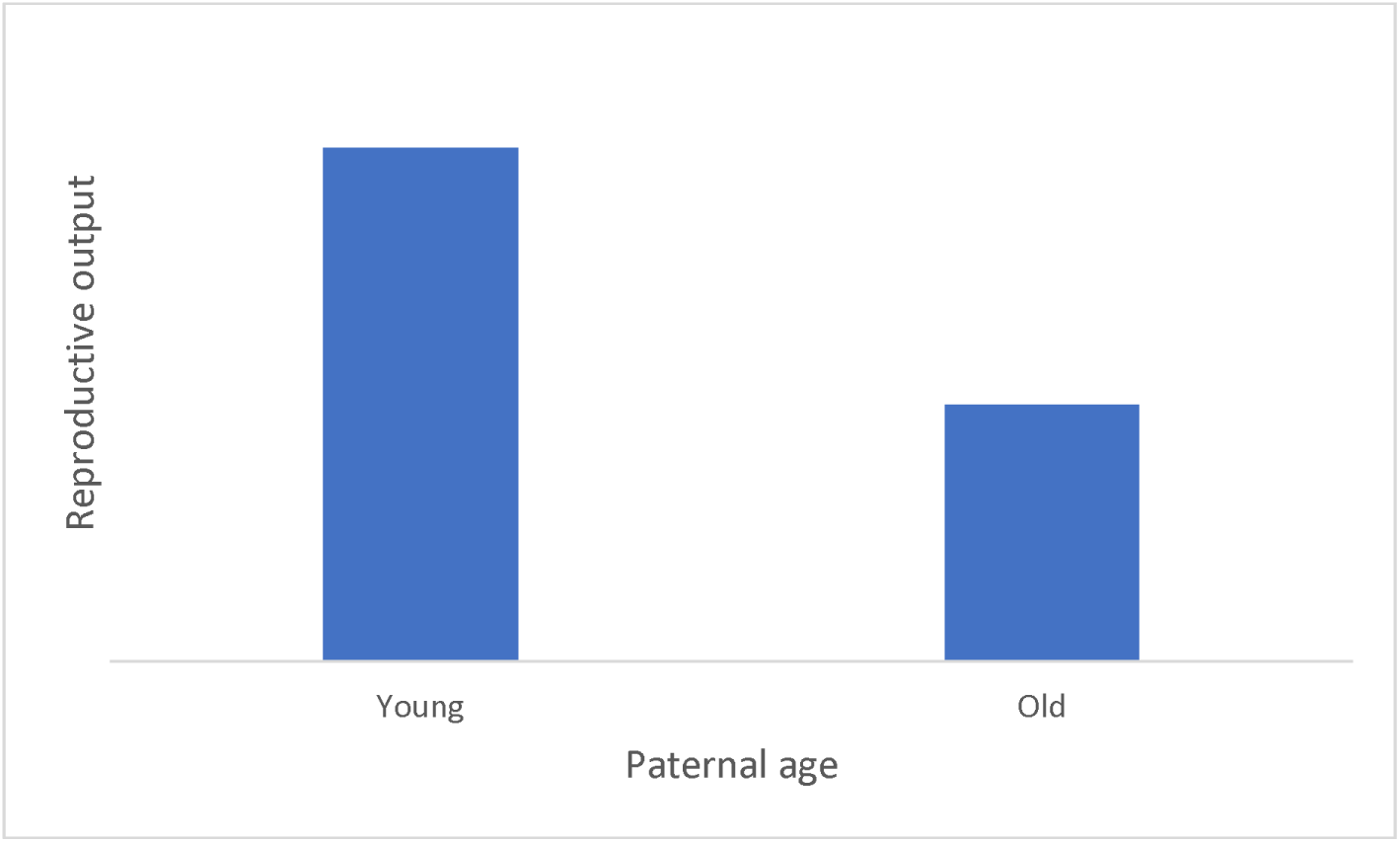
Old males have lower reproductive output than young males due to reproductive senescence
b. Hypothesis 2B: Selective disappearance Prediction: Males who produce fewer offspring will selectively disappear (i.e. die) with age, leading to population level increases in reproductive output with advancing male age. Younger age groups will contain males of poor and high reproductive output while older age groups will contain males of only higher reproductive output, thus have lower variance in reproductive output. **Figure I:**
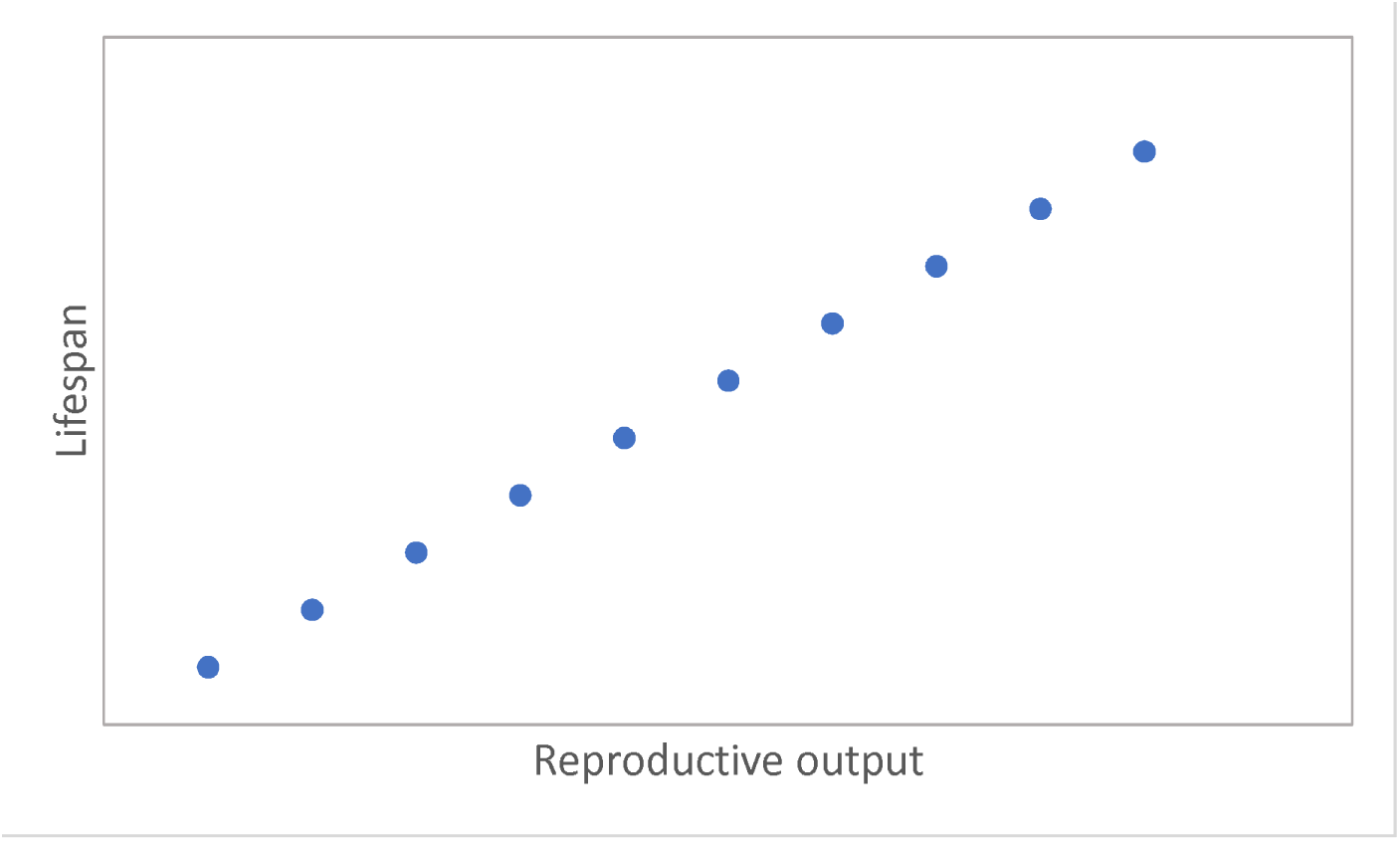
Reproductive output and lifespan co-vary positively due to positive pleiotropy between life-history traits

**Figure J:**
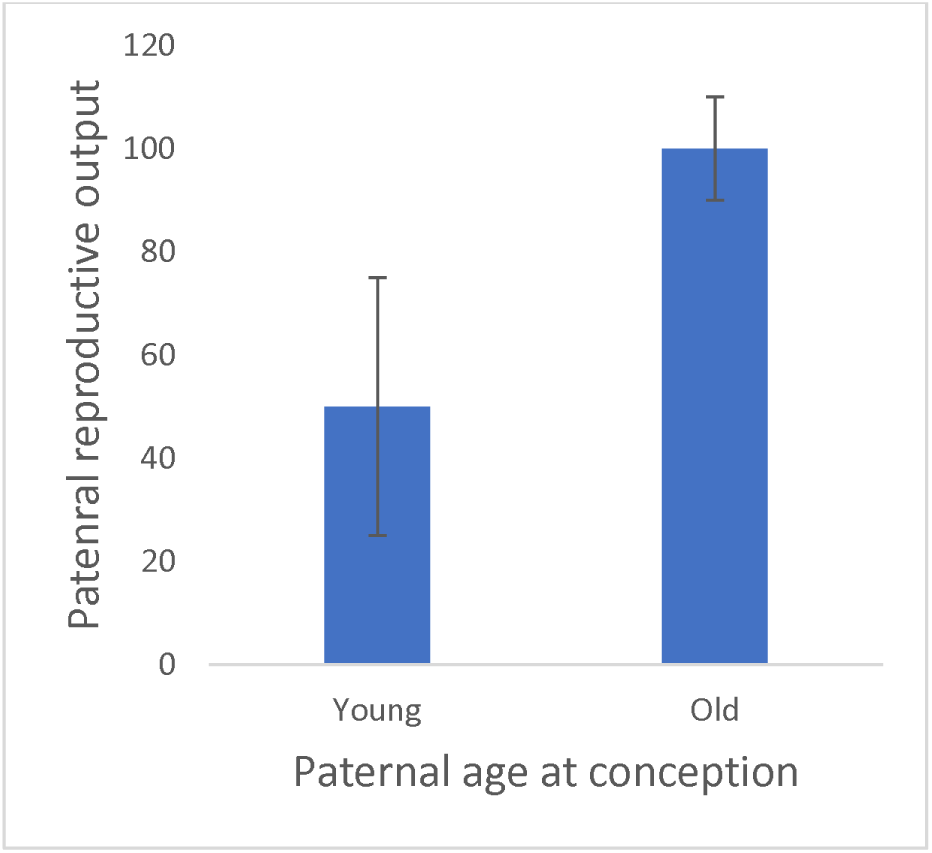
Positive covariances between reproductive output and survival leads to selective disappearance of males and old males having higher reproductive output but lower variance in reproductive output, than young males
c. Hypothesis 2C: Terminal investment Prediction: Older males who are close to dying should invest most in reproduction than young males, or than old males who are not close to dying. **Figure K:**
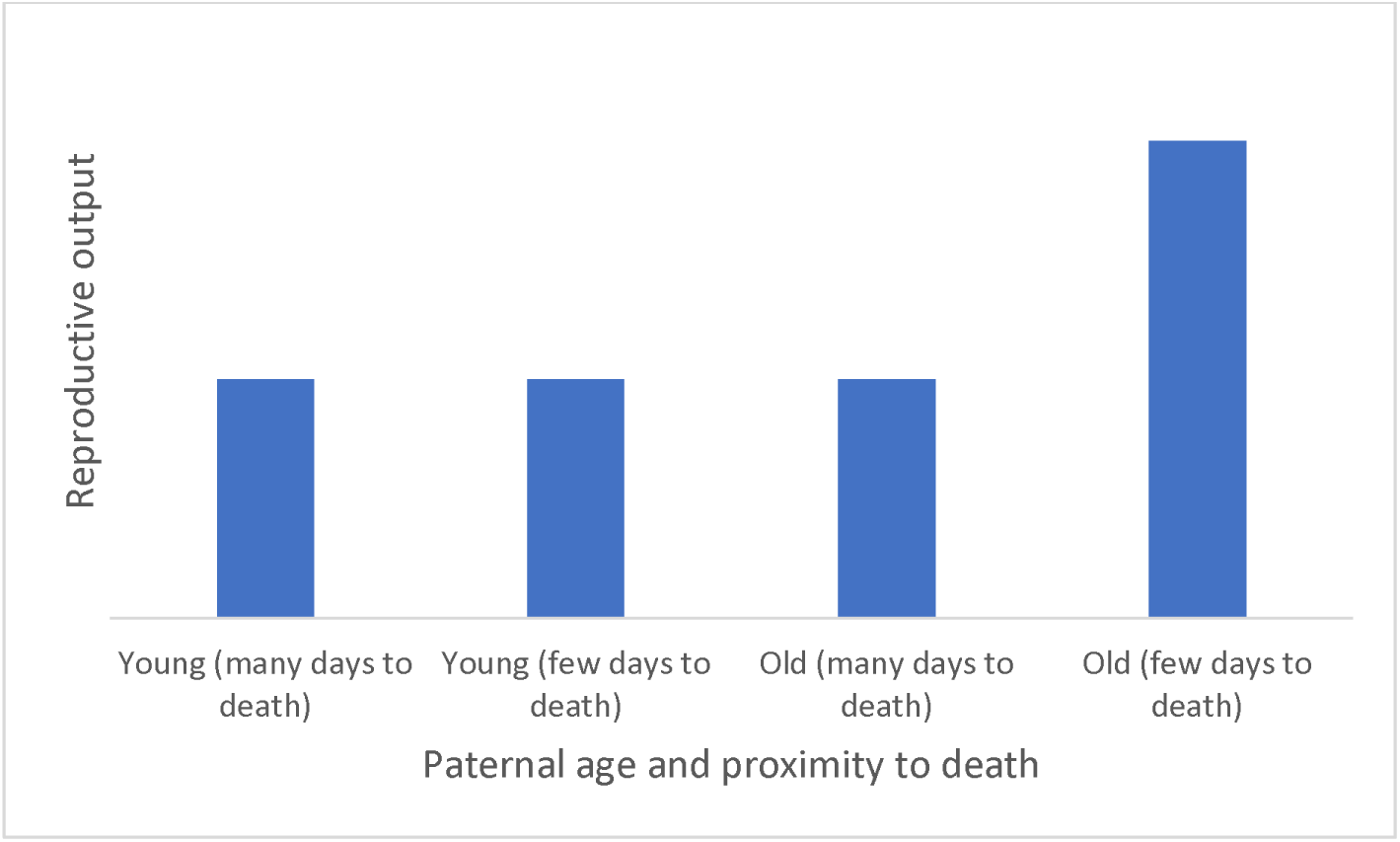
Old males who are about to die soon invest terminally in reproductive output, and have higher reproductive output than old males who are not about to die soon, or than young males.
3. Aim 3:

a. Hypothesis 3A: Lansing effect Prediction 1: Old fathers produce sons with shorter lifespans **Figure L:**
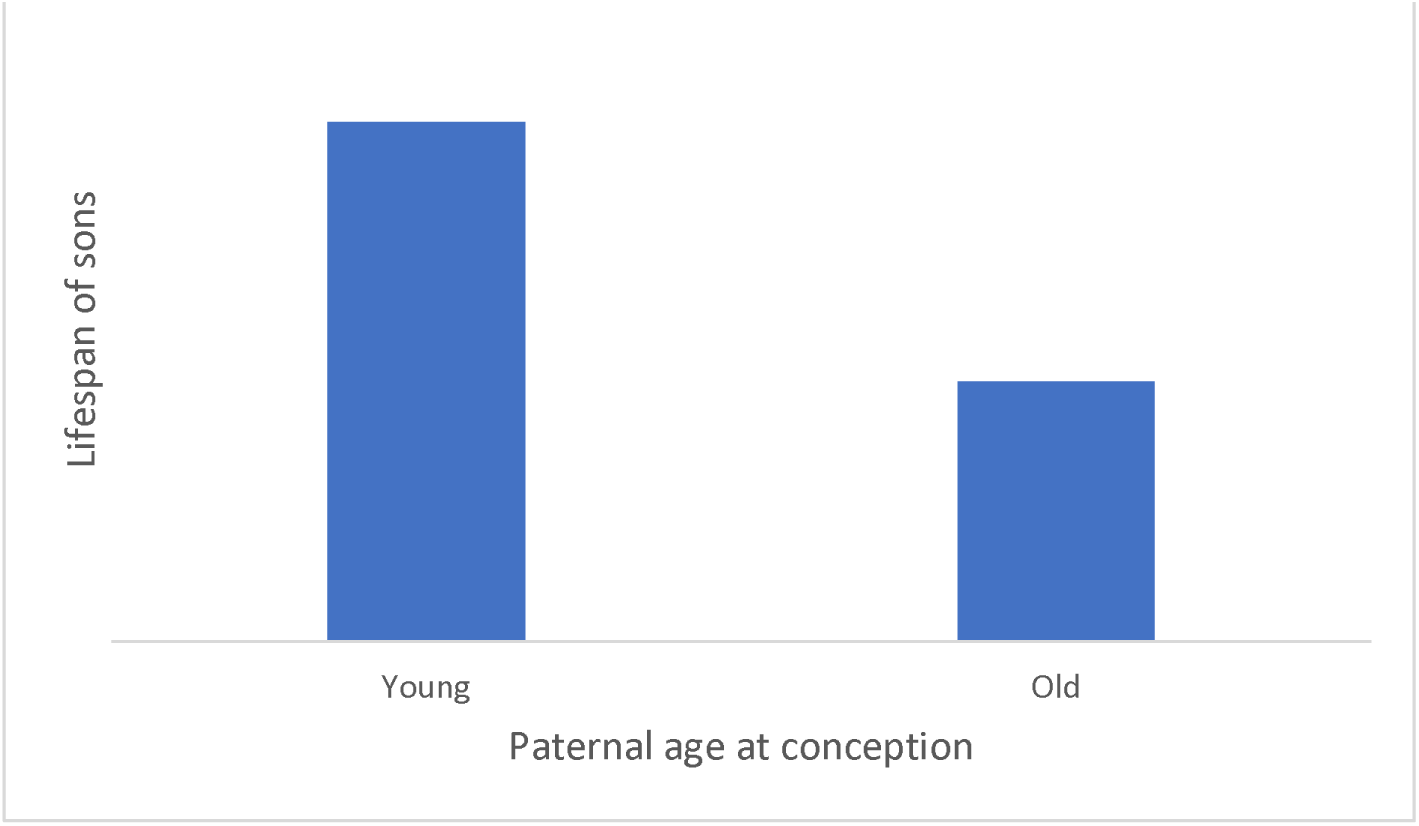
Old father produce sons with lower lifespans due to Lansing effect
b. Hypothesis 3B: Viability selection Prediction: Fathers who mate at older ages are on average longer lived (however have lower variances in lifespan), and produce sons with longer lifespans (but lower variances in lifespan). **Figure M:**
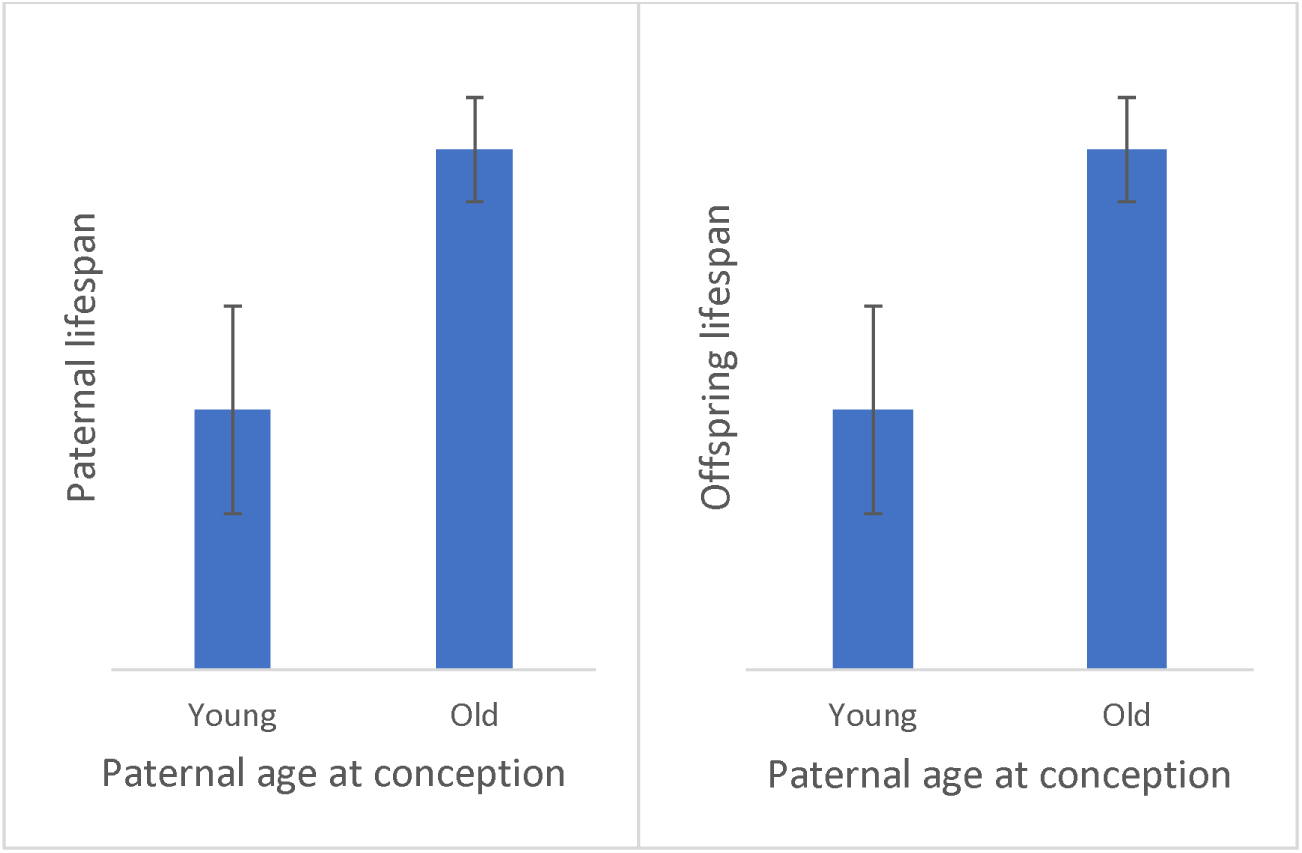
Old fathers have longer lifespans but lower variance in lifespans (also corresponding to H1A), and produce sons with longer lifespans but lower variance in lifespans, than young fathers.

## Appendix 3: Tables and model outputs

**Table S1:** Sample sizes for number of males who were mated at a particular age, and from these, the number of males who produced offspring. Note that at age 60 days, a higher number of males were selected to mate because of the high rate of infertility of old males. Additionally, at 74 days of age, very few males were surviving in the unmated stock population, thus the low sample size.

| PAC (days) | N mated | N produced offspring |
| --- | --- | --- |
| 4 | 25 | 25 |
| 18 | 25 | 23 |
| 32 | 26 | 25 |
| 46 | 26 | 15 |
| 60 | 37 | 14 |
| 74 | 5 | 2 |

**Table S2:** Model output for effects of paternal age at conception (PAC) on probability of surviving three days after mating, with paternal offspring production (F1_count) included as a covariate. Terms highlighted in grey used for interpretation of interaction and main effects.

|  | Estimate | SE | z | P |
| --- | --- | --- | --- | --- |
| (Intercept) | 7.726 | 2.176 | 3.551 | <0.001 |
| PAC | -0.131 | 0.037 | -3.544 | <0.001 |
| F1_count | 0.018 | 0.010 | 1.764 | 0.078 |

**Table S3:** Hurdle model to test interactive effects of paternal age at conception and paternal lifespan, on reproductive output of fathers. Zero inflation model tests effects on the probability of not producing an offspring. Conditional model tests effects on the number of offspring produced, when only data on fathers that produced an offspring were analysed. PAC = paternal age at conception, Paternal_LS = paternal lifespan. Terms highlighted in grey used for interpretation of interaction and main effects.

| Dispersion parameter | 6.19 |  |  |  |
| --- | --- | --- | --- | --- |
| Conditional model (>0) | Estimate | SE | z | P |
| (Intercept) | 3.928 | 0.284 | 13.819 | <0.001 |
| PAC | 0.011 | 0.011 | 1.021 | 0.307 |
| Paternal_LS | 0.001 | 0.004 | 0.126 | 0.899 |
| I(PAC^2) | 0.000 | 0.000 | -3.870 | <0.001 |
| Latency | 0.000 | 0.001 | -0.092 | 0.927 |
| Copulation duration | 0.006 | 0.007 | 0.951 | 0.342 |
| PAC:Paternal_LS | 0.000 | 0.000 | 0.562 | 0.574 |
| Zero-inflation model (1 vs 0) |  |  |  |  |
|  | Estimate | SE | z | P |
| (Intercept) | -9.046 | 5.040 | -1.795 | 0.073 |
| PAC | 0.349 | 0.129 | 2.703 | 0.007 |
| Paternal_LS | 0.094 | 0.071 | 1.312 | 0.190 |
| I(PAC^2) | 0.000 | 0.001 | 0.270 | 0.787 |
| Latency | -0.029 | 0.012 | -2.438 | 0.015 |
| Copulation duration | -0.089 | 0.046 | -1.937 | 0.053 |
| PAC:Paternal_LS | -0.004 | 0.002 | -2.336 | **0.020** |

Main-effects model:
| Conditional model (>0) | Estimate | SE | z | P |
| --- | --- | --- | --- | --- |
| (Intercept) | 3.829 | 0.226 | 16.921 | <0.001 |
| Latency | 0.000 | 0.001 | -0.189 | 0.850 |
| Copulation duration | 0.006 | 0.007 | 0.882 | 0.378 |
| Paternal_LS | 0.002 | 0.002 | 0.963 | 0.336 |
| PAC | 0.016 | 0.006 | 2.598 | **0.009** |
| I(PAC^2) | 0.000 | 0.000 | -3.902 | **<0.001** |
| Zero-inflation model (1 vs 0) |  |  |  |  |
|  | Estimate | SE | z | P |
| (Intercept) | -0.621 | 2.335 | -0.266 | 0.790 |
| Latency | -0.023 | 0.010 | -2.229 | 0.026 |
| Copulation duration | -0.101 | 0.043 | -2.340 | 0.019 |
| Paternal_LS | -0.065 | 0.026 | -2.526 | 0.012 |
| PAC | 0.192 | 0.092 | 2.081 | 0.037 |
| I(PAC^2) | -0.001 | 0.001 | -0.993 | 0.321 |

**Table S4:**
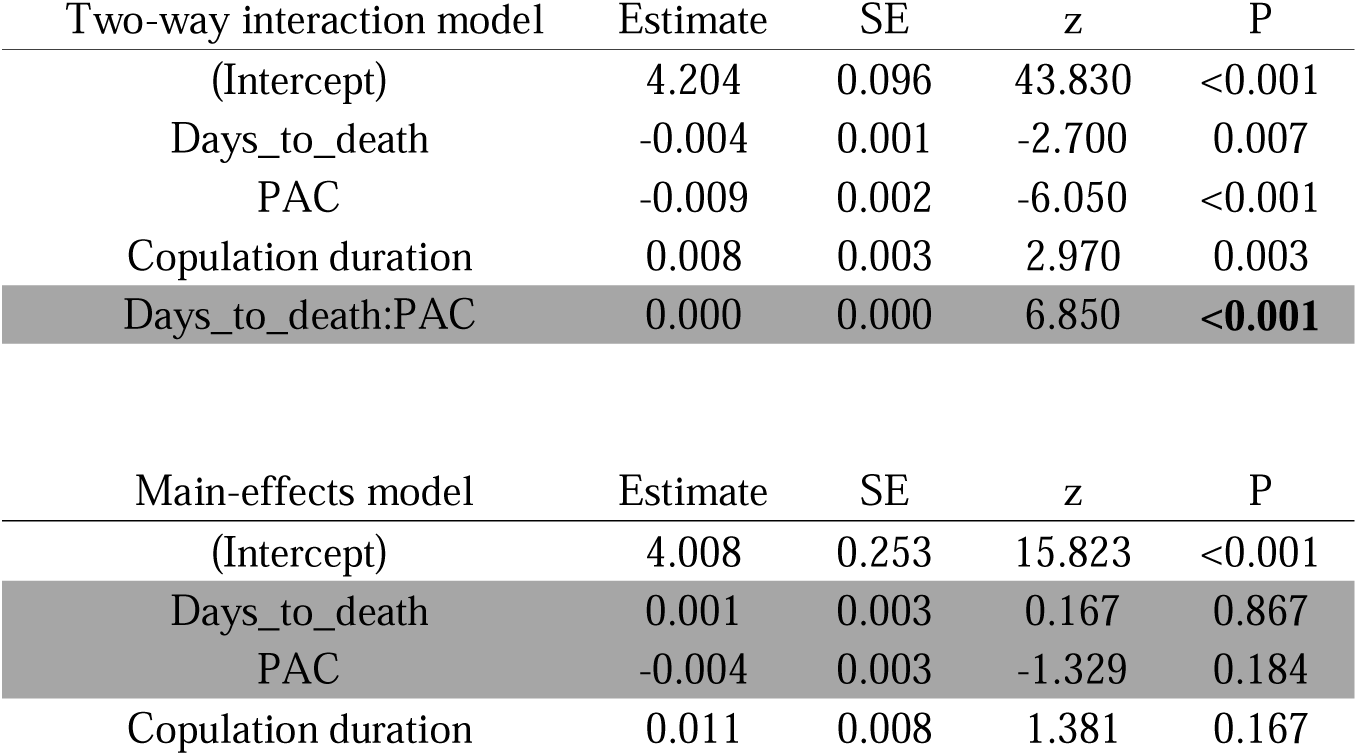
Effects of paternal age at conception and days to death (i.e. time elapsed between death and mating) on the number of offspring produced by fathers. PAC = paternal age at conception. Terms highlighted in grey used for interpretation of interaction and main effects.

| Two-way interaction model | Estimate | SE | z | P |
| --- | --- | --- | --- | --- |
| (Intercept) | 4.204 | 0.096 | 43.830 | <0.001 |
| Days_to_death | -0.004 | 0.001 | -2.700 | 0.007 |
| PAC | -0.009 | 0.002 | -6.050 | <0.001 |
| Copulation duration | 0.008 | 0.003 | 2.970 | 0.003 |
| Days_to_death:PAC | 0.000 | 0.000 | 6.850 | **<0.001** |

| Main-effects model | Estimate | SE | z | P |
| --- | --- | --- | --- | --- |
| (Intercept) | 4.008 | 0.253 | 15.823 | <0.001 |
| Days_to_death | 0.001 | 0.003 | 0.167 | 0.867 |
| PAC | -0.004 | 0.003 | -1.329 | 0.184 |
| Copulation duration | 0.011 | 0.008 | 1.381 | 0.167 |

**Table S5:** Effects of paternal age at conception on lifespans of sons, without accounting for effects of paternal lifespan. Heterogeneous variance function specified as a power function, with parameter showing change in variance in lifespans of sons, with the increasing PAC. PAC = paternal age at conception, F1_count= number of offspring produced by fathers. Terms highlighted in grey used for interpretation of main effects.

| Random effects | SD |
| --- | --- |
| Paternal ID | 4.659 |
| Residual | 22.002 |
| Heterogenous variance parameter (power) | -0.24169 |

| Fixed effects | Estimate | SE | DF | t | P |
| --- | --- | --- | --- | --- | --- |
| (Intercept) | 55.904 | 3.409 | 188.000 | 16.399 | <0.001 |
| F1_count | 0.002 | 0.043 | 98.000 | 0.054 | 0.957 |
| PAC | 0.108 | 0.042 | 98.000 | 2.532 | **0.013** |

**Table S6:** Model output from path analysis (structural equation model), showing the direct and indirect effects (via paternal lifespan) of paternal age at conception, on lifespans of sons. F1_LS = lifespans of sons, PAC = paternal age at conception, Paternal_LS = paternal lifespan, F1_count = number of offspring produced by fathers. Terms highlighted in grey used for interpretation of effects.

|  | Estimate | SE | z | P |
| --- | --- | --- | --- | --- |
| F1_LS ~ PAC (c) | 0.094 | 0.039 | 2.443 | **0.015** |
| Paternal_LS ~ PAC (a) | 0.171 | 0.037 | 4.662 | **<0.001** |
| F1_LS ~ Paternal_LS (b) | 0.115 | 0.060 | 1.920 | 0.055 |

| Covariances: | Estimate | SE | z | P |
| --- | --- | --- | --- | --- |
| Paternal_LS ~~ F1_count | 3.907 | 12.102 | 0.323 | 0.747 |
| F1_LS ~~ F1_count | -0.718 | 12.272 | -0.059 | 0.953 |

| Variances: | Estimate | SE | z | P |
| --- | --- | --- | --- | --- |
| F1_LS | 149.855 | 12.466 | 12.021 | <0.001 |
| Paternal_LS | 145.645 | 12.116 | 12.021 | <0.001 |
| F1_count | 290.526 | 24.169 | 12.021 | <0.001 |

| Defined Parameters | Estimate | SE | z | P |
| --- | --- | --- | --- | --- |
| a*b (indirect effect) | 0.020 | 0.011 | 1.776 | 0.076 |
| c + a*b (total effect) | 0.114 | 0.037 | 3.040 | **0.002** |

### Appendix 4: Supplementary figures

**Figure S1:**
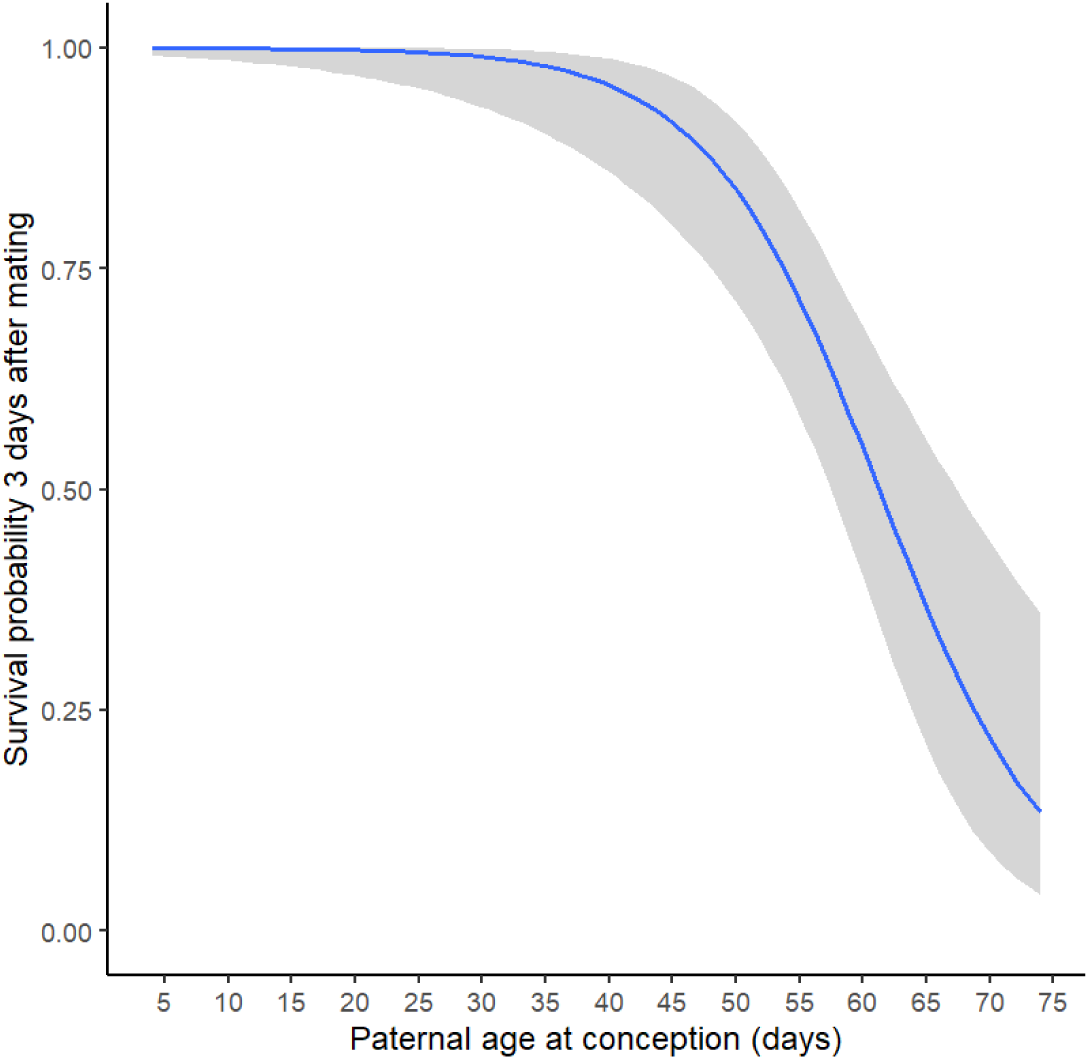
Lower proportion of males are alive 3 days after mating, in older PAC treatments than younger PAC treatments. Shaded areas represent 95% C.I.

**Figure S2:**
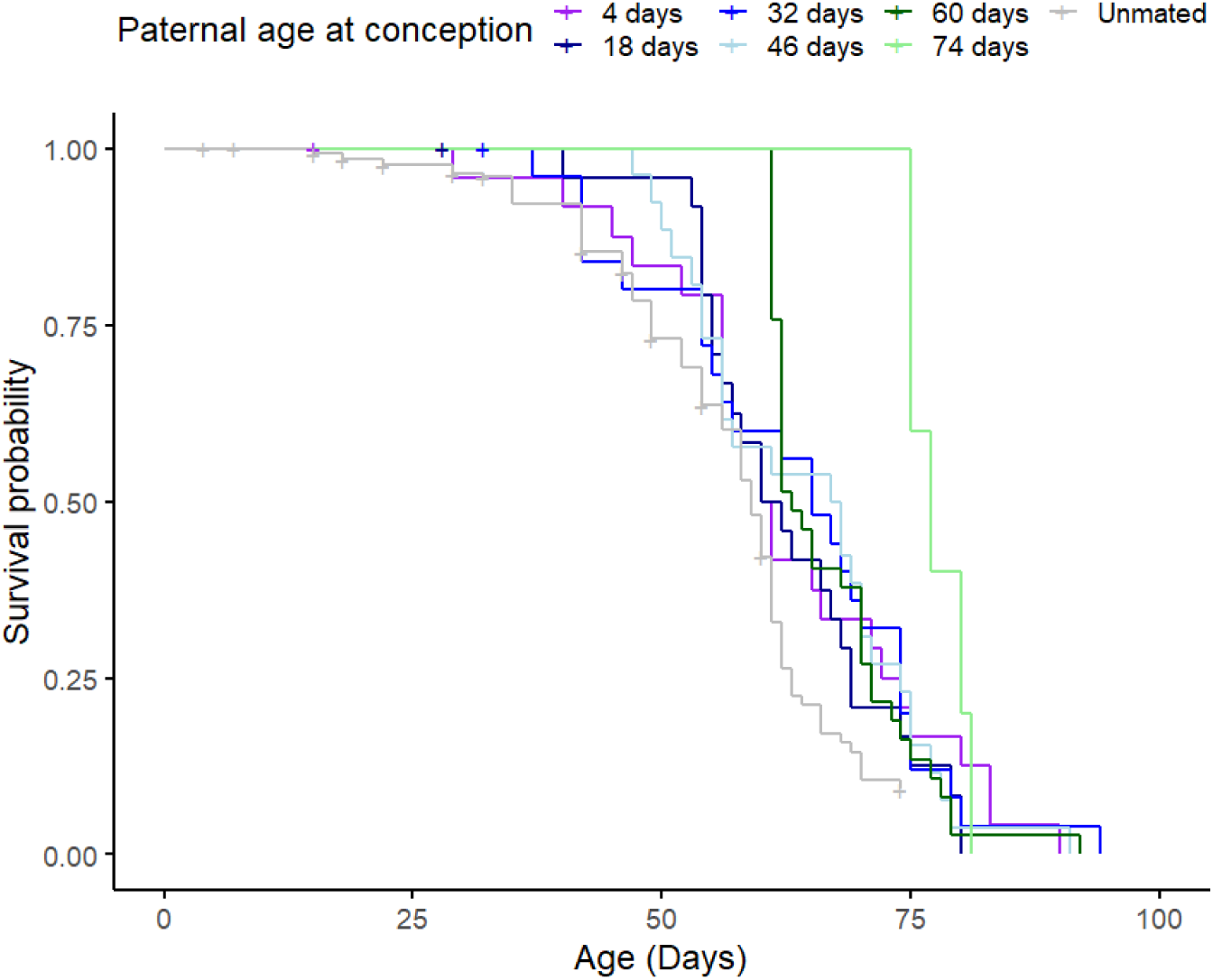
Survival probability of fathers mated at different ages, and males from the unmated experimental population. Fathers who mate at an older age have a lower survival probability at a given age than father who mate at a younger age. “+” signs show censored males.

**Figure S3:**
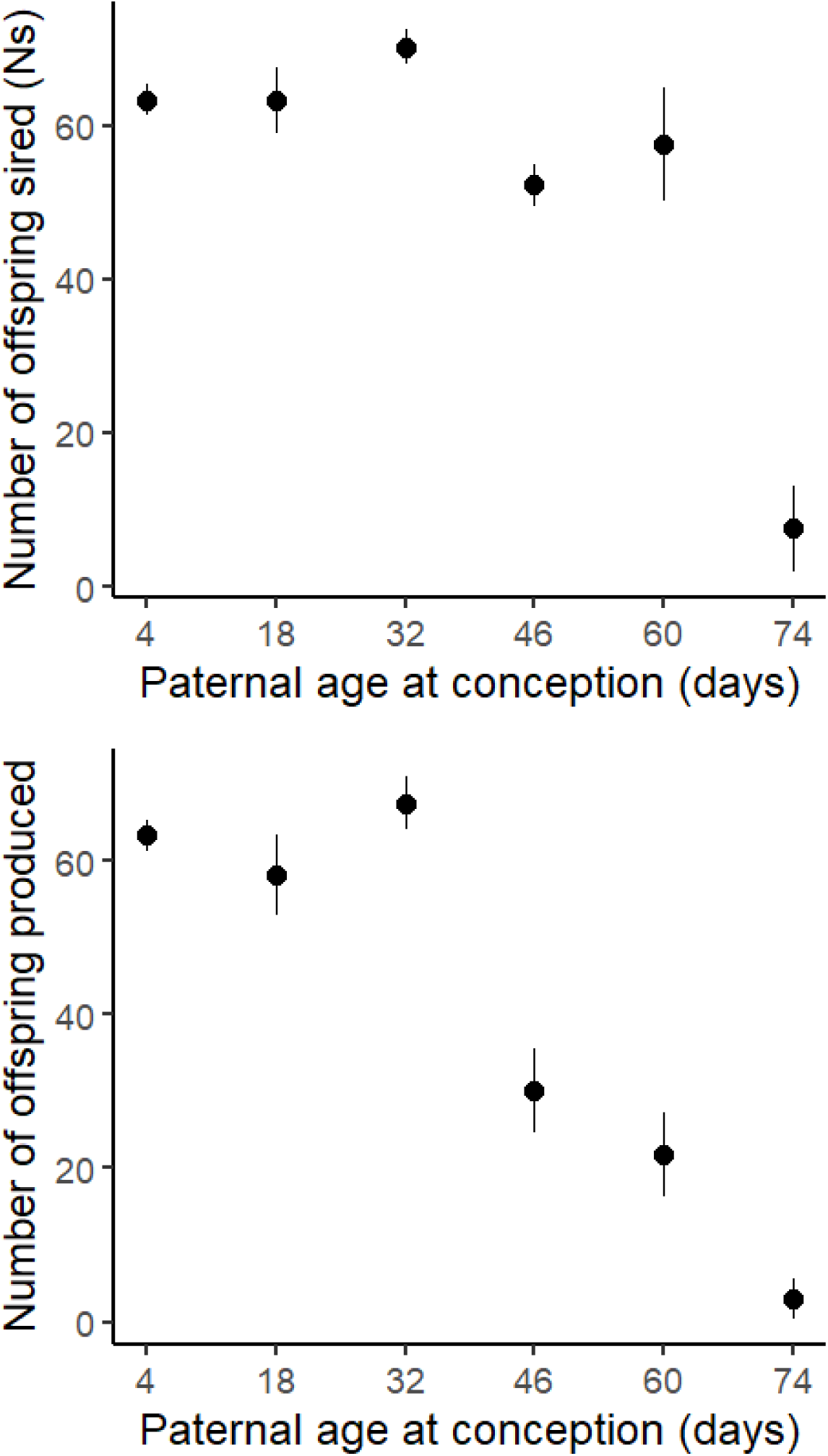
(Top) Paternal age at conception affected the number of offspring sired (excluding fathers who did not produce offspring) in a quadratic way. However, this effect was likely driven by data from PAC of 74 days being lower than other PAC data (also see Figure S4). (Bottom): Effects of paternal at conception on number of offspring produced (including fathers who produced zero offspring). Means and SE shown.

**Figure S4:**
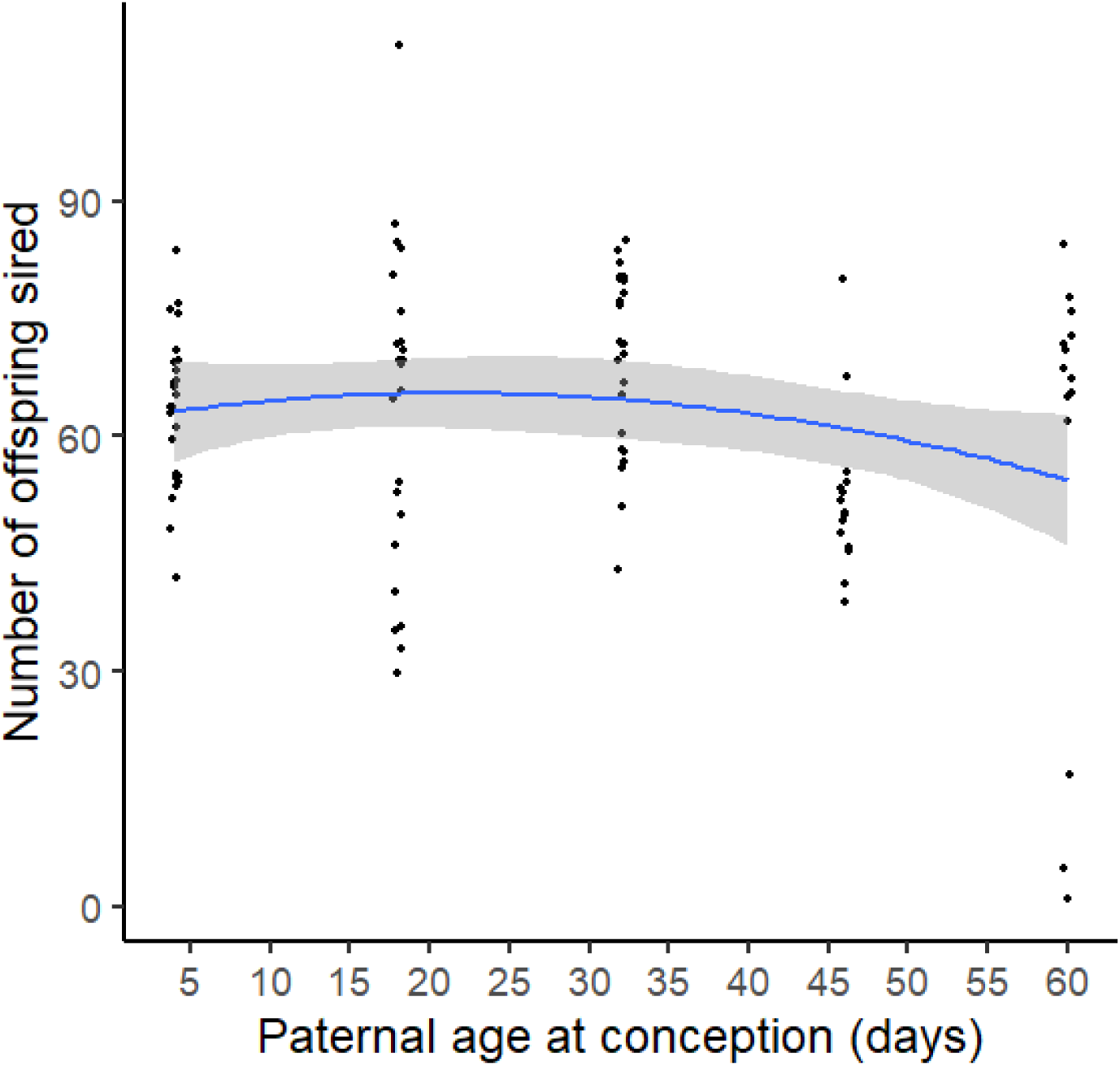
Paternal age at conception affected the number of offspring sired (after excluding fathers who did not produce offspring) in a quadratic way (even when PAC group of 74 days was excluded). However, excluding data from PAC of age 74 days led to a shallow shape of the quadratic curve compared to when this data was included (i.e. Figure 4B). Shaded areas represent 95% C.I.

**Figure S5:**
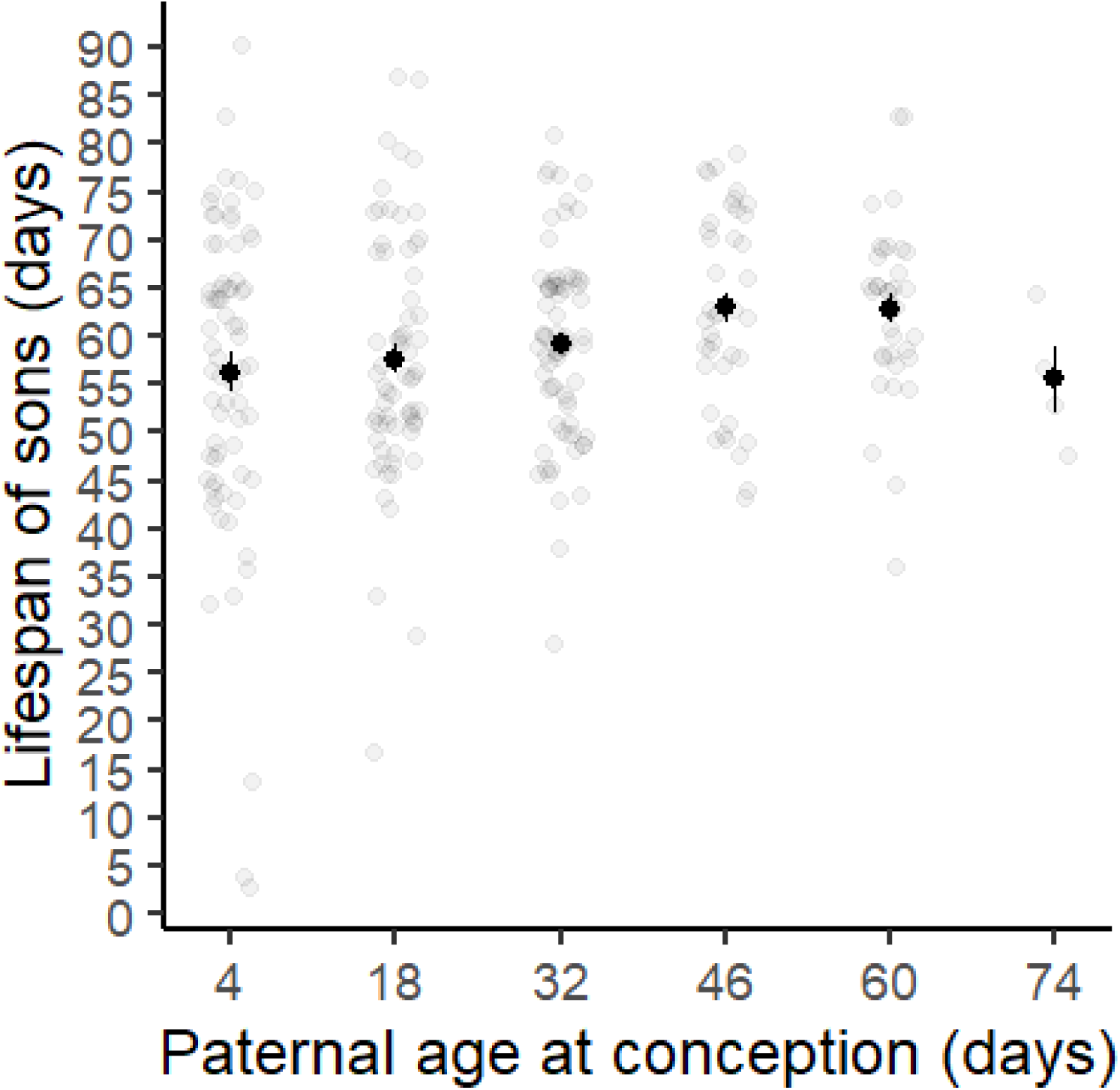
Advancing paternal age at conception increased lifespan of offspring. Means and SE presented

**Figure S6:**
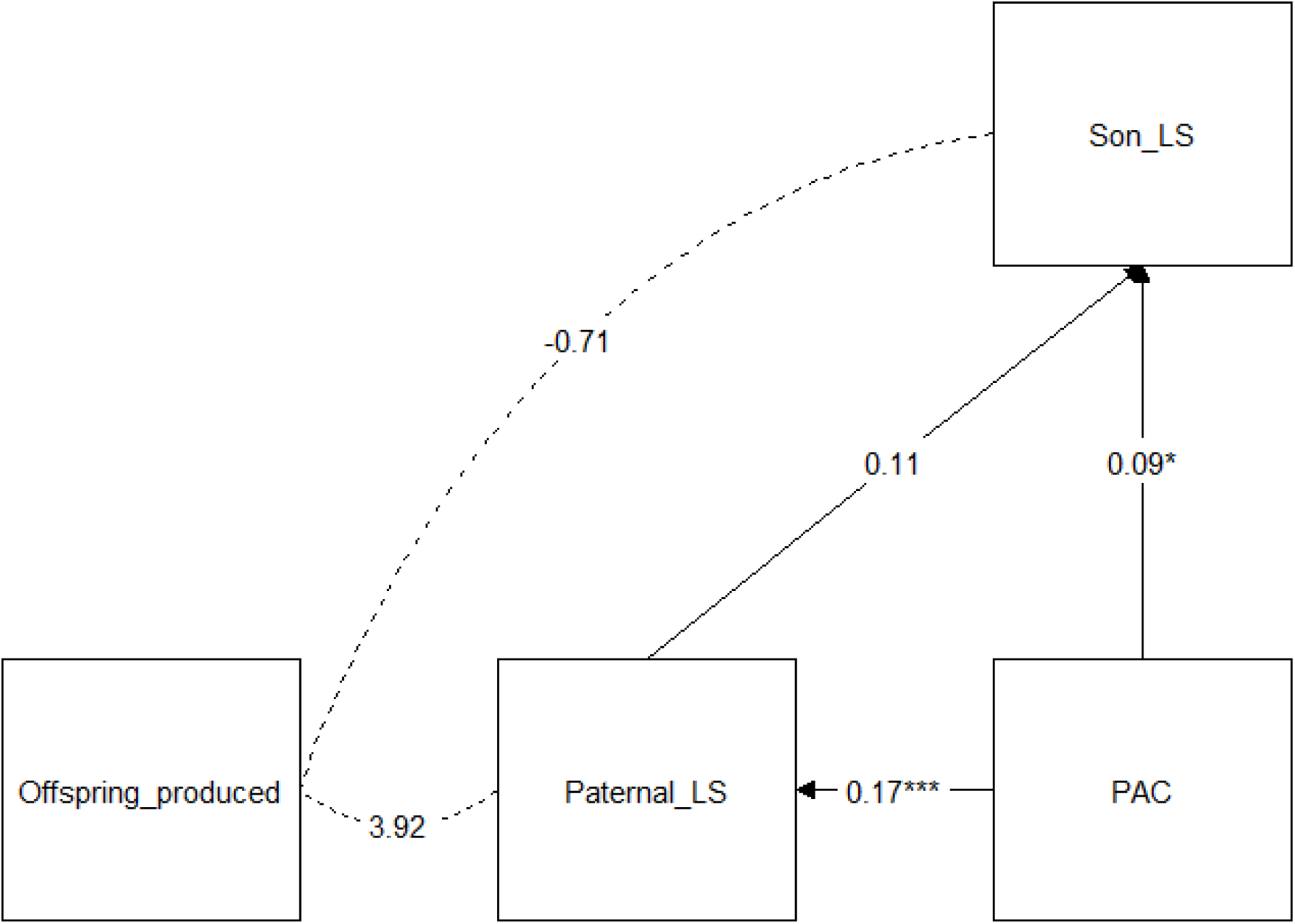
Path analysis showing direct effects of paternal age at conception, and indirect effects via paternal lifespan, on sons’ lifespans, along with covariances (dotted) of paternal lifespan, lifespans of sons, and number of offspring produced. Values show estimates of models, and asterisks represent level of significance.

